# Probing the dissociation pathway of a kinetically labile transthyretin mutant

**DOI:** 10.1101/2023.06.21.545798

**Authors:** Xun Sun, James A. Ferguson, Benjamin I. Leach, Robyn L. Stanfield, H. Jane Dyson, Peter E. Wright

## Abstract

Aggregation of transthyretin (TTR) is associated with devastating TTR amyloid disease. Amyloidosis begins with dissociation of the native tetramer to form a monomeric intermediate that assembles into pathogenic aggregates. This process is accelerated in vitro at low pH, but the dissociation and reassembly of TTR at neutral pH remains poorly understood, due to the low population of intermediates. We use NMR studies with a highly sensitive ^19^F probe that allows deconvolution of relative populations of a destabilized A25T mutant at concentrations as low as 2 µM. The A25T mutation, located at the weak dimer interface, perturbs both the weak and strong dimer interfaces. A tetramer-dimer-monomer (TDM) equilibrium model is proposed to account for concentration- and temperature-dependent population changes. All thermodynamic and kinetic parameters and activation energetics for dissociation of the native A25T tetramer, as well as a destabilized alternative tetramer (T*) with a mispacked F87 side chain, were extracted by van’t Hoff and ^19^F NMR line-shape analysis. The conversion from T to T*, the slowest first-order kinetic step, shows anti-Arrhenius behavior. The ^19^F and methyl chemical shifts of probes close to the strong dimer interface in the dimer and T* species are degenerate, implicating interfacial perturbation as a common structural feature of these intermediate species. Molecular dynamics (MD) simulations further suggest more frequent F87 ring flipping on the nanoscale timescale in the A25T dimer than in the tetramer. Our integrated approach offers quantitative insights into the energy landscape of the dissociation pathway of TTR at neutral pH.

## Introduction

Transthyretin is a blood protein most commonly associated with the transport of thyroid hormone and retinol. Transthyretin amyloidosis is caused by deposition of pathogenic TTR aggregates in tissues and organs including the heart and central nervous system.*^1^* The TTR aggregation pathway begins with dissociation of a native tetramer to form an aggregation-prone monomeric intermediate,*^2^* a process that is accelerated at mildly acidic pH.*^3^* However, wild-type (WT) TTR, as well as most naturally occurring TTR variants, remains predominately tetrameric at physiological concentrations in the blood*^4, 5^* (protomer concentration ~7–21 µM*^6^*). In cerebrospinal fluid (CSF), the TTR concentration range is lower, 0.16–1.6 µM,*^6^* favoring tetramer dissociation for destabilized mutants. For example, a monomer population of 17% has been observed at 298 K and pH 7.0 for an A81T variant at 1.25 µM, but no trace of monomer was seen for the wild type TTR control under the same conditions.*^5^* To study how TTR dissociates at neutral pH, detergents or denaturants have been used to visualize a dimeric species by polyacrylamide gel electrophoresis (PAGE)*^7^* or size exclusion chromatography.*^8^* Dissociation of the tetramer mediated by a strong dimer intermediate has also been studied by molecular dynamics (MD) simulations.*^9, 10^* However, direct experimental observation of lowly populated dissociative intermediates in solutions that are free of detergent or denaturant has not been possible for WT TTR at the physiological concentrations of blood or CSF. Elucidation of the mechanism of TTR dissociation and aggregation at physiological pH is of central importance for understanding potential pathways that result in TTR amyloidosis and disease.

The tetramer formed by the A25T variant is kinetically unstable and is one of the most rapidly dissociating TTR variants reported to date.*^11^* The A25T mutation is located at the weak dimer interface (Figure 1A), where mutations are known to destabilize the native tetramer.*^12^* A25T, a rare variant, aggregates aggressively in CSF and is associated with TTR amyloidosis in the central nervous system.*^11, 13^* In the present work, we used ^19^F NMR to probe the dissociation equilibria of the A25T-C10S-S85C mutant, labeled with 3-bromo-1,1,1-trifluoroacetone (BTFA, the labeled construct is denoted as A25T^F^), over a range of temperatures at neutral pH. The simplest model to account for the ^19^F-NMR data is a tetramer ↔ dimer ↔ monomer (TDM) equilibrium. Global van’t Hoff analysis offers key insights into the thermodynamics of the dissociation pathway of A25T at neutral pH. The analysis revealed the presence of a minor, destabilized form of the tetramer, previously denoted as T*,*^14^* that is more prone to dissociation than the native tetramer. Combining ^19^F-NMR saturation transfer and line-shape analysis, we determined all the rate constants and activation energy barriers of the two-step TDM dissociation pathway. Population analyses using ^19^F and ^13^C-HMQC NMR reveal that the chemical shifts of three probes for the T* and D states are degenerate, indicating similar probe environments in these states. All-atom MD simulations further suggest that both the strong and weak dimer interfaces are perturbed in the A25T dimer. Our work provides quantitative and holistic insights into the thermodynamics and kinetics of the TTR dissociation pathway at neutral pH. These results give important insights into the mechanism whereby the FDA-approved drug tafamidis treats TTR amyloidosis, suppressing TTR dissociation by binding and stabilizing the native tetramer.

**Figure 1.**
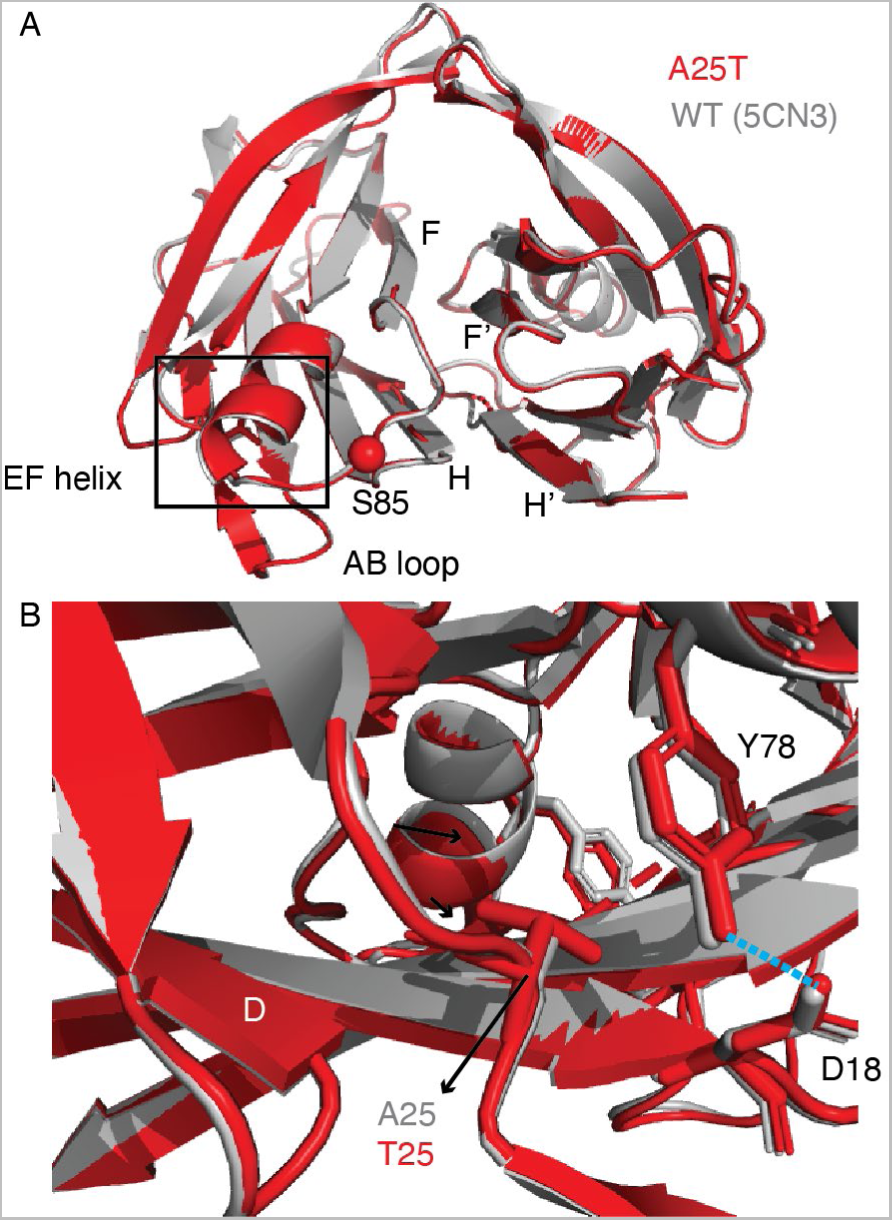
Comparison of the X-ray structures of WT (gray, PDB 5CN3*^15^*) and A25T (red, PDB 8T5X, this work). A. Strong dimer structure present in the asymmetric unit of the A25T structure. Labels show motifs involved in forming the strong dimer interface (F and H strands) and the AB loop that forms a part of the weak dimer interface. The location of the attachment of the ^19^F spin label at position 85 is indicated by a red sphere. B. Expanded view showing residue 25 in the AB loop and the critical D18-Y78 hydrogen bond (blue dashed line) that stabilizes interactions between the AB loop and the nearby EF helix. The short D strand and the EF helix are also labeled.

## Results

### X-ray Crystal Structure of A25T

The A25T mutant was crystallized in space group P 2_1_2_1_2 and diffraction data were collected to a resolution of 1.63 Å. The strong dimer pair of two neighboring protomers was observed in the asymmetric unit (Table S1). Overall, the A25T structure is very similar to a typical WT TTR X-ray structure (with a Cα RMSD = 0.3 Å to PDB 5CN3,*^15^* Figure 1A). The AB loop (residues 18 to 28, including the mutation site) and the nearby EF helix take on a similar conformation in both the A25T and the WT structures (Figure 1B). There is no noticeable change in the crucial hydrogen bond between Y78 hydroxyl and D18 carboxylic oxygen that stabilizes TTR: the hydrogen bond distance of 2.6Å is the same in both structures. As shown by over 200 TTR X-ray structures*^16^* and a previously published A25T X-ray structure (PDB 3TFB,*^13^* Figure S1), X-ray structures of TTR seldom if ever reveal pronounced conformational perturbations caused by individual mutations.

### Solution ^15^N-NMR spectroscopy of A25T

Solution NMR using ^15^N and ^13^C-labeled protein can give a better idea of the structural and dynamic changes occurring as a consequence of mutations. Several amide and Cα cross peaks are broadened or missing from 2D and 3D triple resonance spectra of A25T. Overlaid ^1^H, ^15^N-TROSY spectra of A25T and WT TTR are shown in Figure 2A. Residues in the AB loop, D strand or EF helix are broadened or shifted substantially in the A25T spectrum relative to WT, and V93 and V94 in the strong dimer interface are also broadened (Figure 2A). A number of resonances were completely missing from 3D triple resonance spectra (trHNCA and trHN(CO)CA) of A25T. Residues without visible amide or Cα connectivities are mapped onto the A25T structure in Figure 2B. These residues are mostly located in the AB loop and the DAGH sheet (Figure 2B). The broadening also propagates to the F strand (residues 91 to 97), which forms part of the strong dimer interface (Figure 2C). Both the weak and, to a lesser degree, the strong dimer interfaces are perturbed by the A25T mutation. For backbone amides that can be assigned, chemical shift perturbations (CSP) are observed for residues extending from the N-terminus of the EF helix (S77) to the C-terminus of the E strand (V71, Figure 2D). Interestingly, Y78 has similar chemical shifts in A25T and WT, consistent with the similarity of the Y78 (side chain)-D18 hydrogen bond distances and orientations in their X-ray structures (Figure 1B). The secondary Cα chemical shifts of A25T are consistent with the secondary structures observed in the A25T X-ray structure (Figure S2).

**Figure 2.**
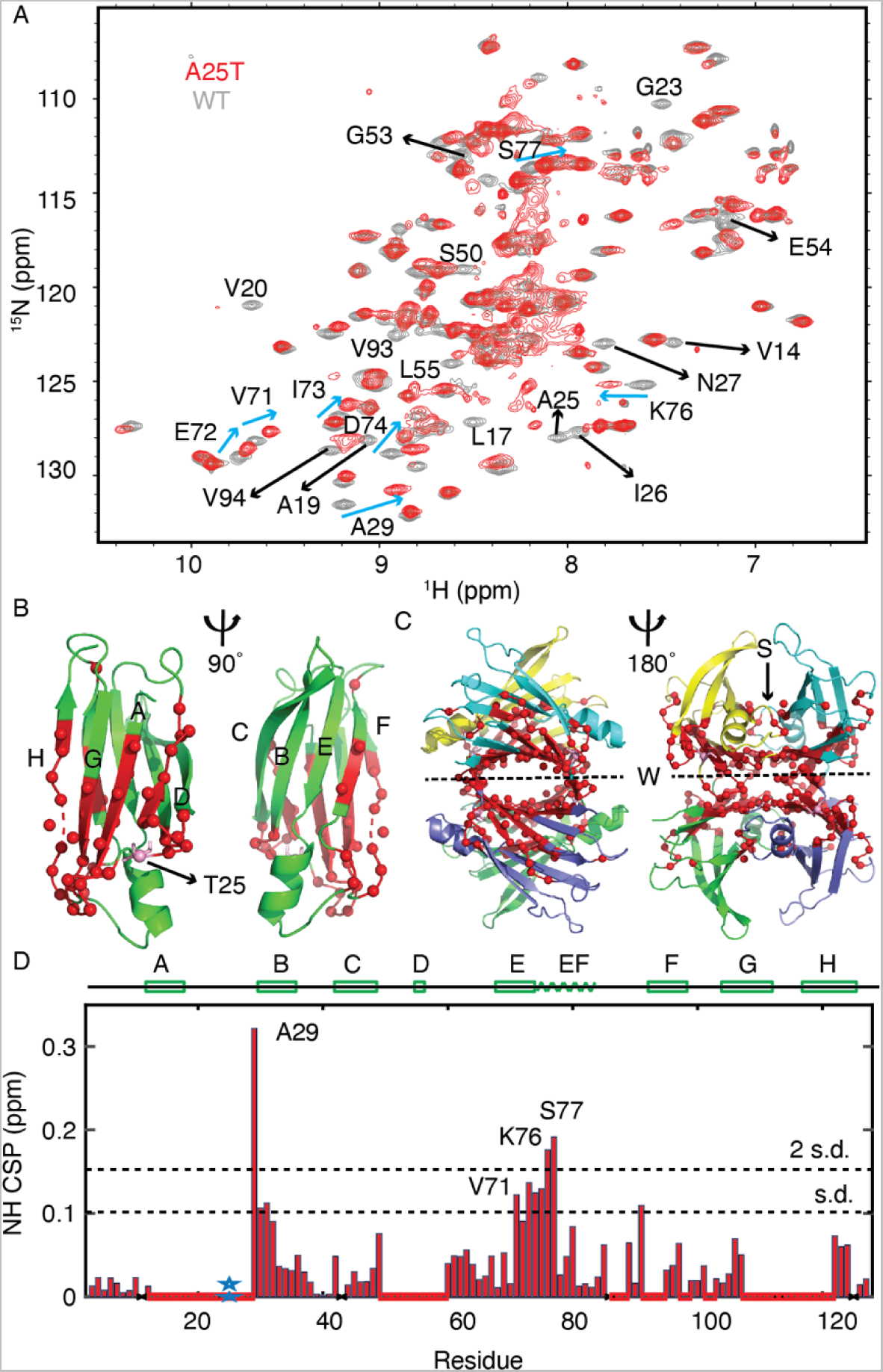
Perturbation of the NMR spectrum by the A25T mutation. A. Superposition of ^15^N-TROSY spectra of A25T (red) and WT (gray) at 298 K and 800 MHz. Representative residues in the AB loop, D strand or EF helix that are broadened or shifted substantially in the A25T spectrum relative to WT are labeled. Broadened residues at the strong dimer interface (V93 and V94) are also labeled. B. Residues without visible amide NH or Cα peaks in 3D trHNCA and trHN(CO)CA spectra are mapped as red spheres on two views of the A25T protomer structure with β-strands labeled. C. Mapping of residues with broadened cross peaks onto the symmetry-imposed tetramer structure of A25T; the majority of these residues are localized at the weak (W) and strong (S) dimer interfaces. D. Chemical shift perturbation CSP = √[(Δδ_H_)^2^ + (Δδ_N_/5)^2^] of A25T versus WT. Dashed lines denote mean+1 or mean+2 standard deviations. Secondary structures are indicated above the figure. Red boxes on the x-axis denote residues with broadened resonances and the blue star indicates the A25T mutation.

### Concentration series monitored by ^19^F-NMR

We then asked whether the perturbed dimer interfaces promote dissociation of the native A25T tetramer. To probe dissociation of A25T at low µM concentration, we introduced a sensitive ^19^F BTFA probe at position 85, using the construct A25T-C10S-S85C-BTFA (denoted as A25T^F^). This fluorine probe senses distinct oligomerization states of TTR.*^17^* At 310 K, pH 7.0 and a concentration of 1.25 µM TTR (corresponding to physiological levels in the CSF), three peaks were observed in the ^19^F spectrum of A25T^F^ (Figure 3A). The main peak at −83.92 ppm arises from the native tetramer (T). The shoulder at −84.01 ppm was initially identified as T*, corresponding to a mispacked tetramer state with reduced stability as described previously.*^14^* Later analysis indicates that this peak also includes the signal from a dimer intermediate (see below). The third peak at −84.20 ppm corresponds to the monomer (M). The same ^19^F chemical shift difference (0.28 ppm) was observed between the T and M peaks of the corresponding WT construct C10S-S85C-BTFA (TTR^F^) at pH 4.4.*^17^* The appearance of the M peak at 1.25 µM is also seen at neutral pH in ^19^F spectra of the K80E and K80D mutants, where the Schellman helix C-capping motif at the EF helix and EF loop is perturbed, but not in TTR^F^ at neutral pH.*^5^* Incubation with the FDA-approved drug tafamidis results in the loss of the shoulder and monomer peaks and an increase in the intensity of the T peak (Figure 3B), consistent with the mode of action of tafamidis in binding at the central hydrophobic cavity to stabilize the native tetramer.*^18^*

**Figure 3.**
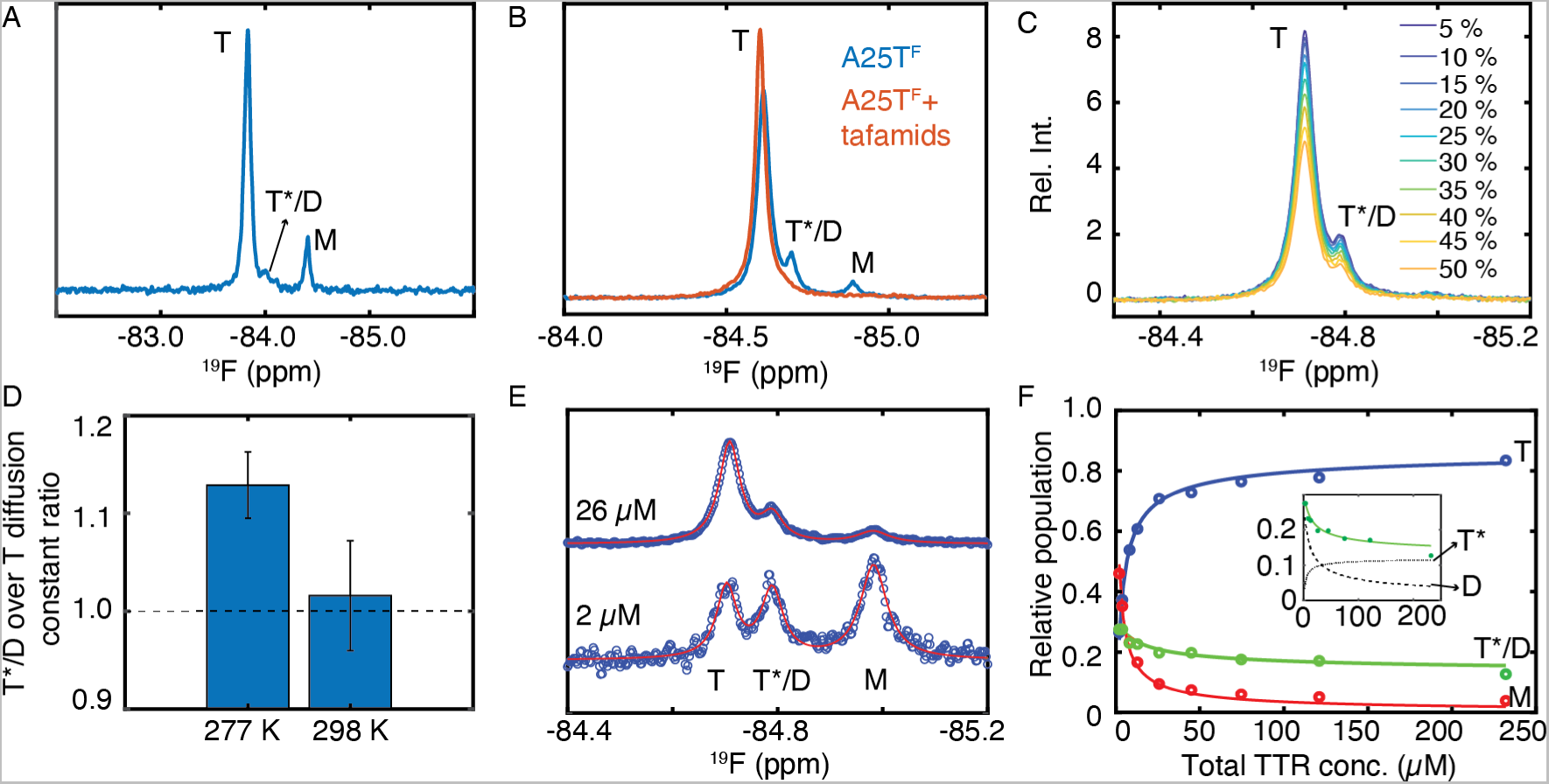
^19^F-NMR spectra of A25T^F^ under physiologically relevant conditions. (A) ^19^F spectrum of A25T^F^ at 1.25 µM collected at 310 K and pH 7.0 with T, T*/D and M peaks labeled. (B) ^19^F spectra of 30 µM A25T^F^ without (blue) and with 50 µM tafamidis (red) at 281 K and pH 7.0. The incubation time was 60 mins. (C) ^19^F-DOSY measurements of 230 µM A25T^F^ at 277 K with 10 z-gradient strengths. (D) The translational diffusion constant ratio of the T*/D peak over the T peak at 277 and 298 K for 230 µM A25T^F^. The results show that a species smaller than tetramer is present under the shoulder peak and is more highly populated at 277 K than at 298 K. The error is propagated from fitting uncertainty from 50 bootstrapped data sets. (E) Concentration dependence of A25T^F^ spectrum at 277 K with 3-state Lorentzian fits shown in solid lines and data as open circles. For clarity, only selected concentrations are shown; full data set fits are shown in Figure S3. (F) Fitting relative populations using the TDM model where the ^19^F chemical shift of T* and D is degenerate as the shoulder peak. Inset: both the T* and D contribute to the observed shoulder peak. The model fits are shown in solid lines. Alternative models with worse fitting statistics are shown in Figure S4.

To probe the oligomerization state of the shoulder peak, we performed ^19^F DOSY experiments for A25T^F^ at 277 K where the population of the shoulder peak is enhanced compared to higher temperatures (Figure 3C). The relative translational diffusion constant of the shoulder peak over that of the T peak is 1.13 ± 0.03, which is reduced to 1.02 ± 0.03 at 298 K (Figure 3D). This indicates that a species that diffuses faster than tetramer is more populated under the shoulder peak at 277 K than 298 K. To further probe the identity of the species under the shoulder peak, we recorded a series of spectra at decreasing A25T^F^ concentration at 277 K (Figure 3E and S3). At the lowest protein concentrations, the proportions of the monomer and shoulder peaks are greatly increased relative to the tetramer peak. The simplest dissociation model to fit the concentration series spectra is a tetramer-dimer-monomer (TDM) model where both the mispacked tetramer T* and the dimer D contribute to the shoulder peak (Figure 3F and Scheme 1).

**Scheme 1.**
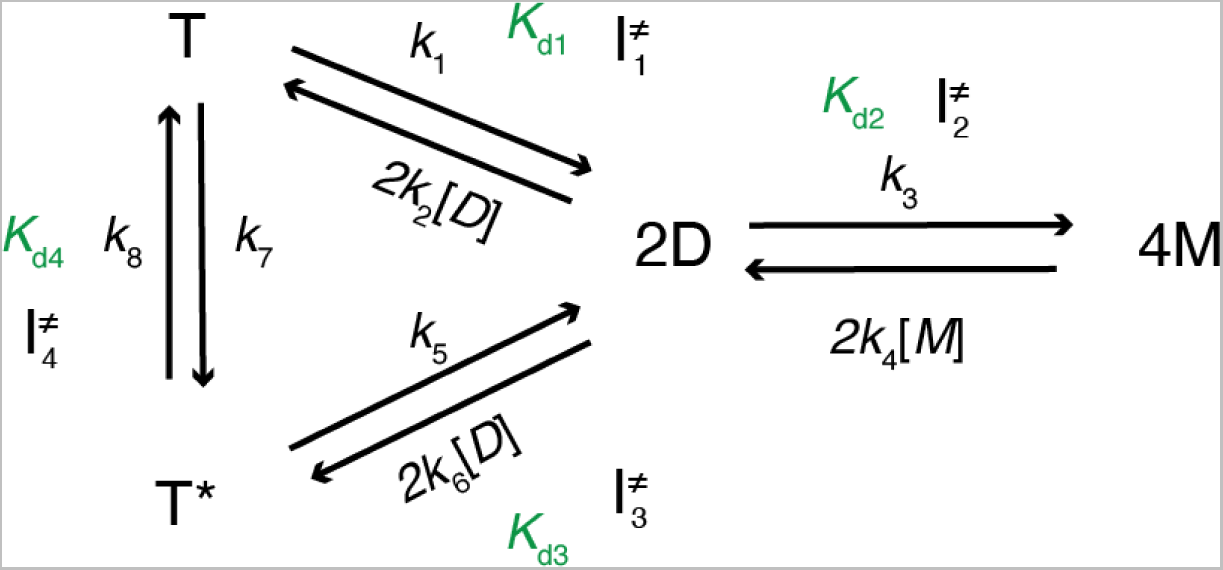
The TDM dissociation model for A25T at pH 7.0. Transition states are labeled as 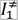 to 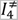

Alternative models without D or that invoke a trimer species either give worse fits or are unable to constrain *K*_d_ parameters (Figure S4). Fitting the 277K concentration series to the TDM model yields three *K*_d_ values between 0.5 and 4 µM where the T* and D are 7-fold and 8-fold more dissociation-prone than the native T species, respectively (Table S2). According to this model, at low concentrations, the dimer is primarily responsible for the enhanced height of the shoulder peak (Figure 3F inset). The similar ^19^F chemical shifts of T* and D suggest comparable environments for the S85C-BTFA probe, which is located in the EF loop near the strong dimer interface (Figure 1A). This finding also suggests that the D intermediate species is likely to correspond to the strong dimer.*^19^* If the ^19^F D peak represented the weak dimer with a disengaged F-F’ and H-H’ strong dimer interface (Figure 1A), the environment around the S85C-BTFA probe would be more similar to the M species than to T*, which is inconsistent with the distinct ^19^F chemical shifts of the M and the D species in A25T^F^. A comparison of the behavior of WT, A25T and the monomeric F87A mutant by SDS PAGE analysis also shows that the strong dimer interface is largely maintained in A25T^F^ compared to TTR^F^, although a slight reduction of the stability of the strong dimer is observed (Figure S5), consistent with the slight perturbation observed for the F strand residues by ^15^N-TROSY spectroscopy (Figure 2B–C).

### Thermodynamics of the A25T TDM dissociation pathway

We recorded ^19^F NMR spectra of A25T^F^ at 6, 30 and 230 µM concentration over a range of 8 temperatures from 277 to 298 K and quantified the relative populations of the T, T*/D, and M peaks using 3-state Lorentzian fits (Figure 4A–D and S6). The observed population changes are not due to changes in ^19^F *T*_1_ values, which are around 0.3 s for all species at 298 K or 277 K (Table S3). A global fit of the temperature titration data sets to the van’t Hoff equation gives *K*_d_ values for each of the three dissociation processes in the TDM model in Scheme 1 that are statistically similar to those obtained by fitting of the concentration series at 277 K alone (Table S2). Additional concentration-dependent data were acquired at 298 K and all five data sets were fitted globally (Figure 4E–F). The fitted thermodynamic parameters are listed in Table 1.

**Figure 4.**
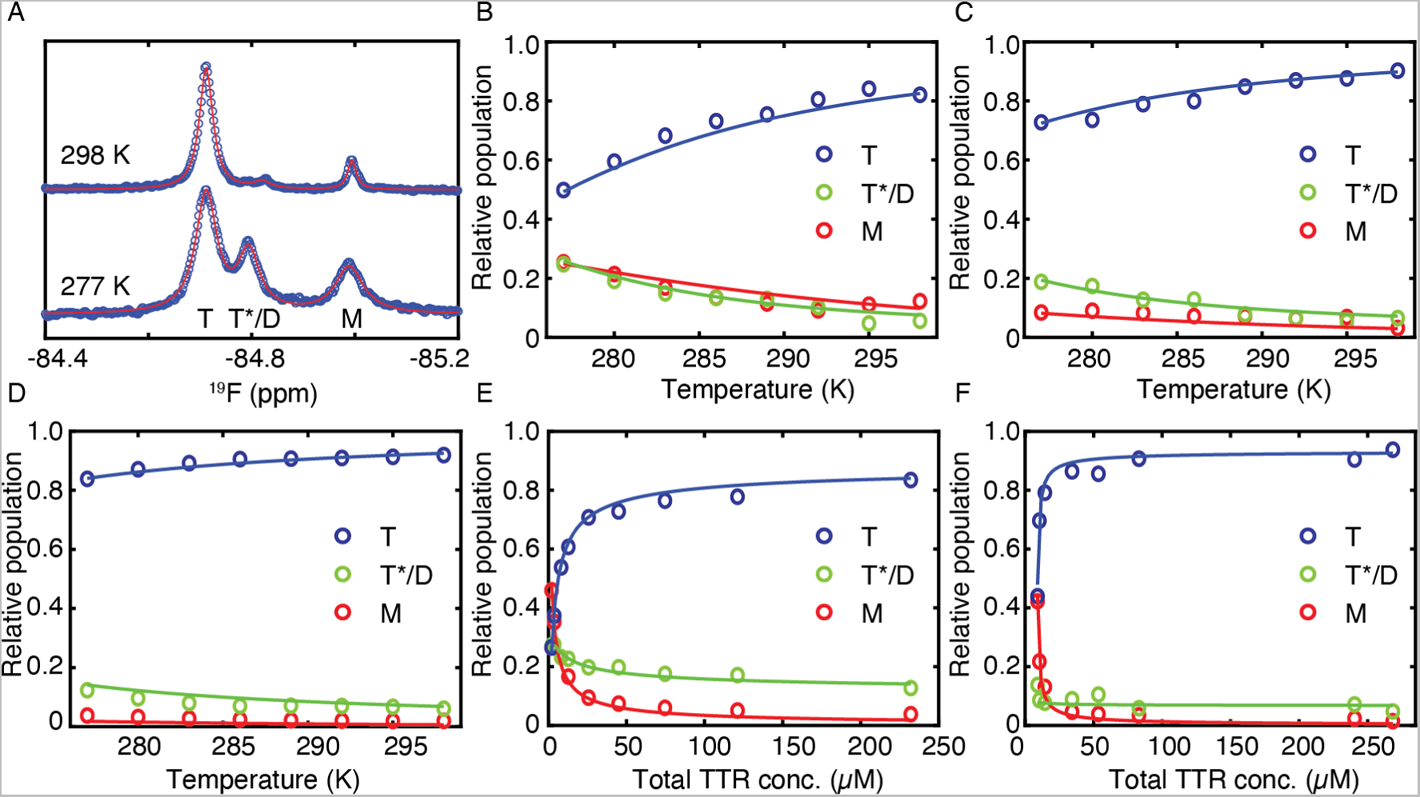
Global van’t Hoff analysis of temperature and concentration series of A25T^F^ data sets. A. Temperature titration of 6 µM A25T^F^ spectra aligned using the T peak in the 277 K spectrum. Data are shown in circles and 3-state Lorentzian fits are in solid lines. For clarity, data from only two temperatures are shown. See fitting results in Table 1 and the full data set fits in Figure S6. B–D. Relative population of T, T*/D, and M states as a function of temperature at A25T^F^ concentrations of 6 µM (B), 30 µM (C), and 230 µM (D). E-F. Changes in relative population of T, T*/D, and M states with changes in concentration at 277 K (E) and 298 K (F). These data sets were used in the van’t Hoff analysis of the TDM dissociation model for A25T^F^. Model fits are shown in solid lines.

**Table 1.** Fitted Δ*H* and Δ*S* parameters by van’t Hoff analysis for the TDM dissociation model of A25T^F^

| Equilibrium<sup>a</sup> | T ↔ 2D | T* ↔ 2D | D ↔ 2M |
| --- | --- | --- | --- |
| $\Delta H$ (kcal/mol) | $\Delta H_1 = -42 \pm 4$ | $\Delta H_3 = -38 \pm 5$ | $\Delta H_2 = 4 \pm 2$ |
| $\Delta S$ (cal/mol/K) | $\Delta S_1 = -179 \pm 15$ | $\Delta S_3 = -160 \pm 16$ | $\Delta S_2 = -10 \pm 8$ |
| $\Delta G$ (kcal/mol) at 277 K | $\Delta G_1 = 8.0 \pm 0.1$ | $\Delta G_3 = 6.8 \pm 0.1$ | $\Delta G_2 = 6.9 \pm 0.1$ |
| $\Delta G$ (kcal/mol) at 298 K | $\Delta G_1 = 11.8 \pm 0.3$ | $\Delta G_3 = 10.2 \pm 0.3$ | $\Delta G_2 = 7.1 \pm 0.2$ |

<sup>a</sup> $\Delta H$ and $\Delta S$ for the forward reaction in each equilibrium (from left to right).

Knowledge of Δ*H* and Δ*S* for each dissociation process allows computation of Δ*G* as a function of temperature (Figure 5A). The Δ*H* and Δ*S* for the formation of T and T* are both positive (Table 1), indicating that this process is mostly driven by hydrophobic interactions. This observation likely reflects the formation of the central hydrophobic channel during tetramerization.*^5, 18^* The increased entropy upon forming tetramer is likely related to the release of hydration water molecules from the weak dimer interface as it becomes buried in the tetramer (Figure 1A). Lowering the temperature from 298 to 277 K weakens *K*_d,1_ for dissociation of T from 2 nM to 0.5 µM (Table S2), providing a thermodynamic rationale for the observed cold denaturation of the TTR tetramer.*^20^* From 277 to 298 K, *K*_d,3_ for dissociation of T* is consistently 8–14 fold weaker than *K*_d,1_ as T* is 1.2–1.6 kcal/mol less stable than T.

**Figure 5.**
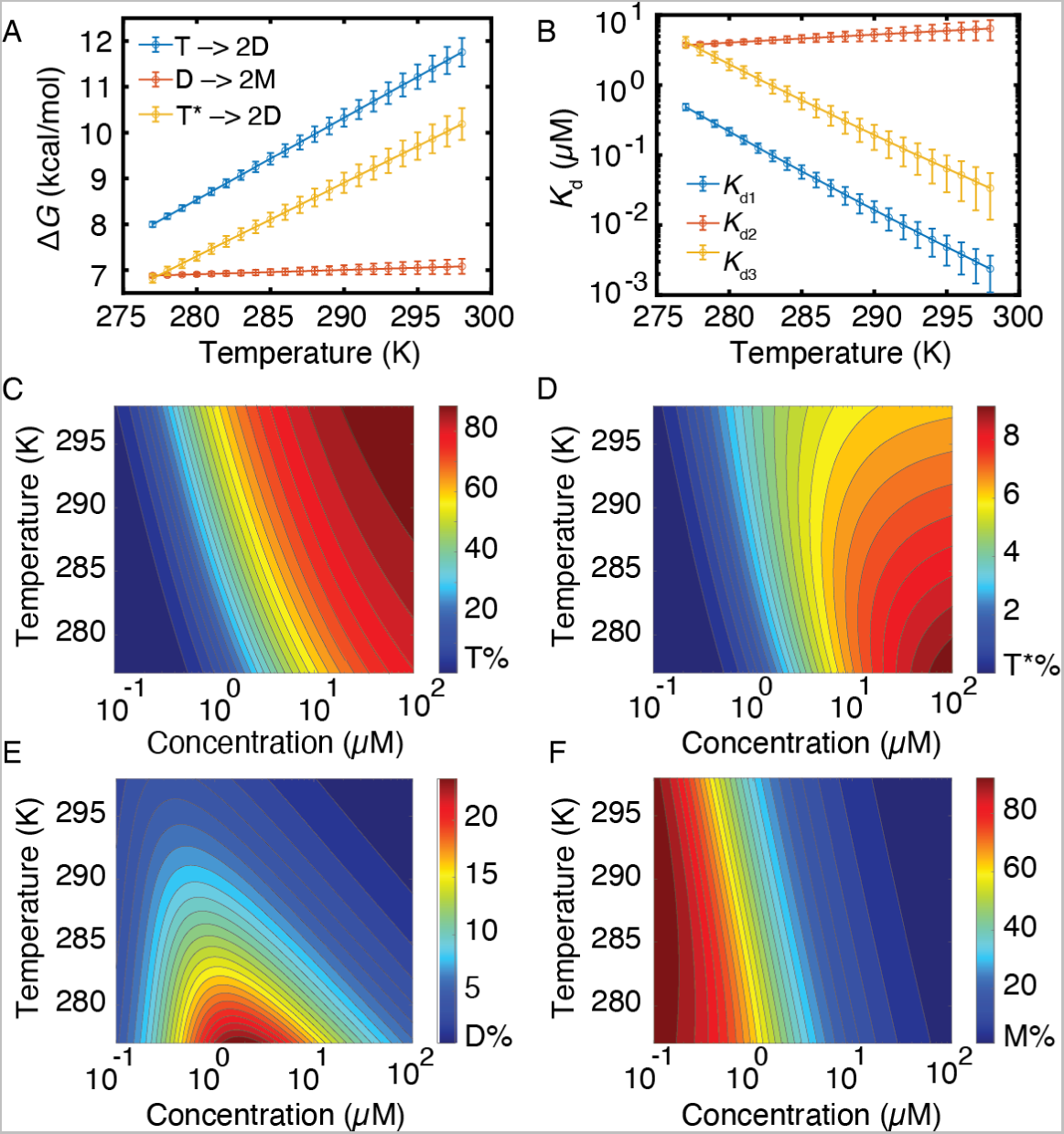
Free energy and population from the van’t Hoff analysis. A. Temperature-dependent changes in free energy for T and T* dissociation to D (blue and orange respectively) and for dissociation of D to M (red). B. Dissociation constants derived from the equilibria in A. Errors are 1 s.d. from 100 bootstrapped datasets. C–F. 2D-contour plots of relative protomer population in the form of T (C), T* (D), D (E) and M (F) of A25T^F^ at pH 7.0 for various temperatures and concentration conditions.

By contrast, *K*_d,2_ for dimer dissociation remains around 4–6 µM between 277 and 298 K (Table S2), a much weaker temperature dependence than *K*_d,1_ or *K*_d,3_ (Figure 5B). This temperature insensitivity arises because the absolute value of Δ*H*_2_ (for dissociation of dimer to form monomer) is about an order of magnitude smaller than Δ*H*_1_ or Δ*H*_3_. The formation of the strong dimer (D) is mainly driven by enthalpy (Δ*H*_2_ = −4.2 kcal/mol), consistent with the extensive hydrogen-bonding between the H and the H’ strands (Figure 1A). The 2-dimensional contour plots of the relative populations of the T, T*, D and M species as a function of temperature and concentration summarize their thermodynamic stabilities (Figure 5C–F). High concentrations favor both T and T* tetramers at the cost of M, and D behaves like a dissociation intermediate. Due to its lower free energy, T is more favored at higher temperature than T*, especially above 10 µM. Within the physiological concentration range (0.1–20 µM), the population of dimer is less than 5% at 298 K, explaining the similar diffusion constants of the T*/D shoulder and T peak (Figure 2D) and the lack of a noticeable dimer peak in the size-exclusion chromatogram (Figure S7).

### Temperature dependent population shifts by ^13^C NMR

The methyl ^1^H-^13^C-HMQC spectrum (Figure 6A) shows five main tetramer cross peaks for the δ1 methyls of the five Ile residues in A25T, which were assigned by mutagenesis (Figure S8A-D). The I26δ1, I68δ1 and I84δ1 cross peaks are more intense than those from I73δ1 and I107δ1. Two additional minor cross peaks are observed at positions similar to those of the I68δ1 and I84δ1 cross peaks in spectra of the monomeric mutant F87E-V122I (Figure S8E), and likely arise from the corresponding Ile residues in a small population of the A25T monomer. Temperature-dependent ^13^C-HMQC spectra confirm that the relative population of the two minor peaks increases at low temperature, where tetramer dissociation is enhanced. However, the population of M predicted from the ^19^F-NMR van’t Hoff analysis (Table 1) underestimates the relative intensities of the minor peaks in the ^13^C-HMQC spectrum. Rather, both the intensity and volume of the minor peaks and their temperature dependence can be accounted for if it is assumed that they arise from all three of the intermediates T*, D, and M (Figure 6B). We infer that the chemical shifts of the I68 and I84 δ1 methyls of T* and D are coincident, as also observed for the ^19^F chemical shift in spectra of A25T^F^ (Figure 3).

**Figure 6.**
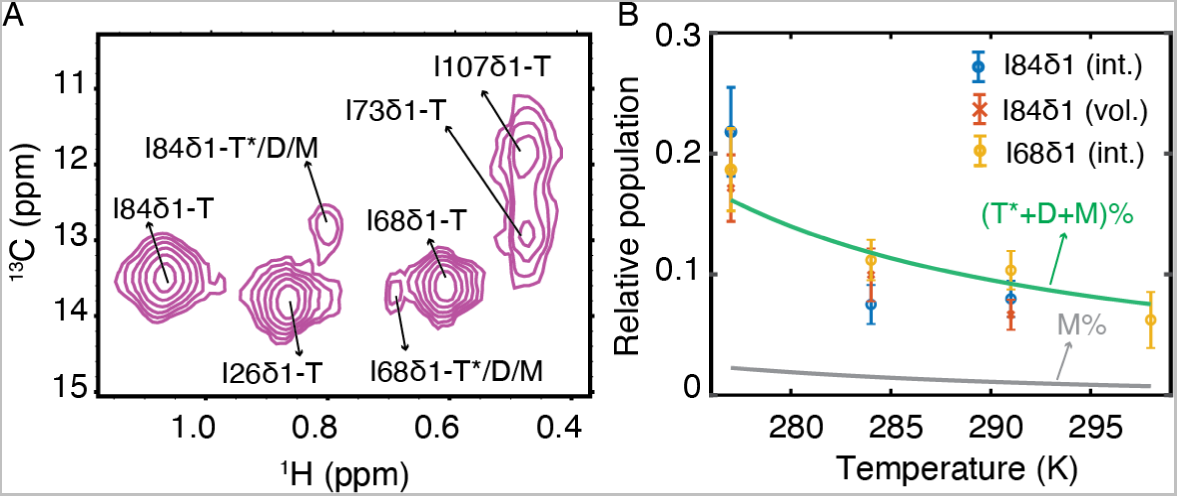
Characterization A25T using ^13^C-HMQC spectroscopy. A. Ile δ1 methyl region in the ^13^C-HMQC spectrum of A25T at 216 µM, 277 K and pH 7.0 with assignments indicated. The T residues were assigned by mutagenesis and the assignments of I68δ1 and I84δ1of the monomer to the minor peaks were made by transfer from the spectrum of monomeric V122I-F87E (Figure S8). B. The M species alone (gray line) cannot account for the relative populations of the minor peaks of I84δ1 and I68δ1 (blue, red and orange markers by intensity or volume as labeled). The population of M and the sum of the T*, D and M populations (green line) were determined from the thermodynamic parameters in Table 1. The population analysis suggests that the methyl chemical shifts of the minor peaks of I84δ1 and I68δ1 are degenerate for T*, D and M.

### Kinetics and activation energetics from ^19^F-NMR line-shape analysis

Rate constants for the TDM model (Scheme 1) were extracted from ^19^F-NMR line-shape analysis of the concentration-dependent experiments at 277 K and 298 K (Figure 4E, F). Based on the four *K*_d_ constraints, we only need to fit one of the two rate constants for each equilibrium step. Since ^19^F chemical shifts for T, T*/D, and M are independent of concentration (Figure S9), peak center positions were constrained to further reduce the number of fitting parameters. The rate constant ratios *k*_3_/*k*_4_ and *k*_7_/*k*_8_ were determined by ^19^F saturation transfer and *K*_d_ constraints (Figure S10 and Table 2) so that the only floating kinetic parameters are *k*_1_ and *k*_5_. In addition, three ^19^F linewidths at 277 K and two linewidths at 298 K were fitted (see Methods). The fitted spectra are shown in Figure 7. The fitted and measured kinetic parameters are summarized in Table 2 and the schematic in Figure 8E, and the extracted linewidths are listed in Table 3. Note that the 2D→T rate constant (~3 × 10^9^ M^−1^ s^−1^) approaches the diffusion limit at 298 K (~7 × 10^9^ M^−1^ s^−^1, estimated by the Smoluchowski coagulation equation*^21^* at 298 K), which has been suggested to be feasible for reactions with the interaction potential greater than 10 *k*_B_*T*.*^22^* For comparison, the absolute free energy change for 2D→T is 11.8 kcal/mol at 298 K (~20 *k*_B_*T*).

**Figure 7.**
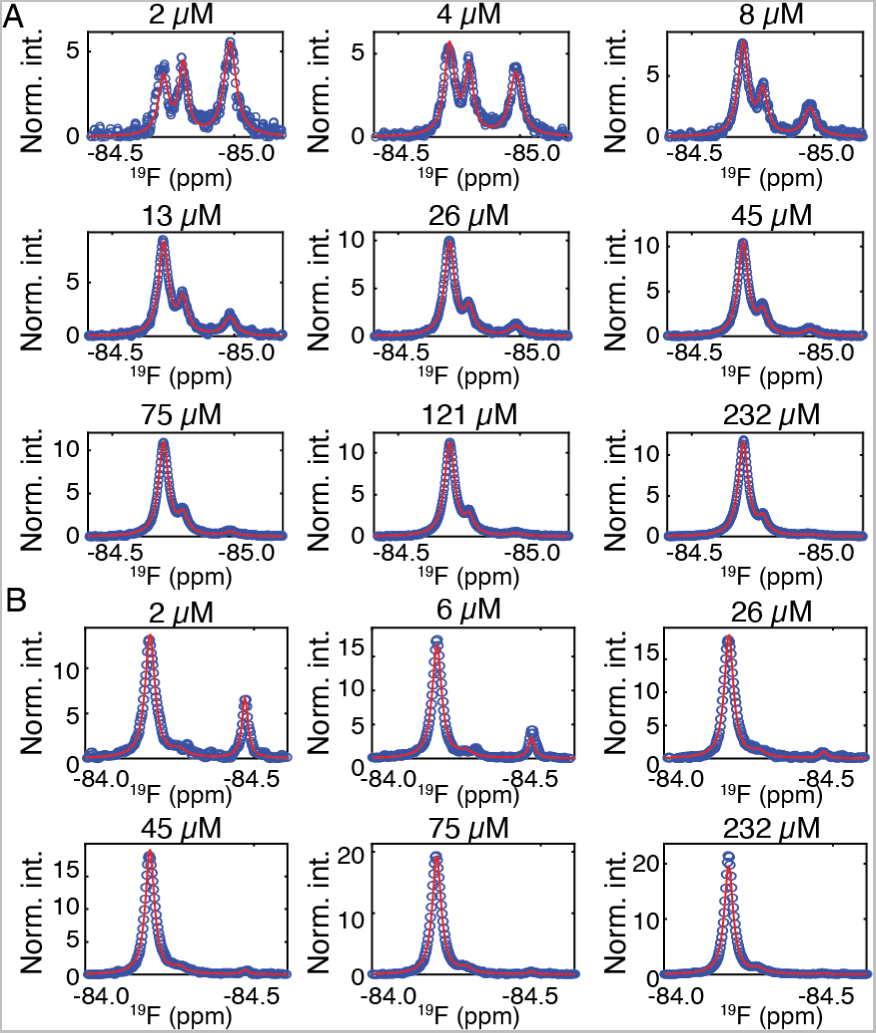
^19^F-NMR line shape analysis for the A25T concentration series. A–B. Concentration-dependent ^19^F spectra at 277 K (A) and 298 K (B) are shown in blue circles and line-shape fits are in red solid lines. The intensity of each spectrum was normalized to ensure the same total peak area for all spectra. Concentrations are shown at the top of each panel.

**Figure 8.**
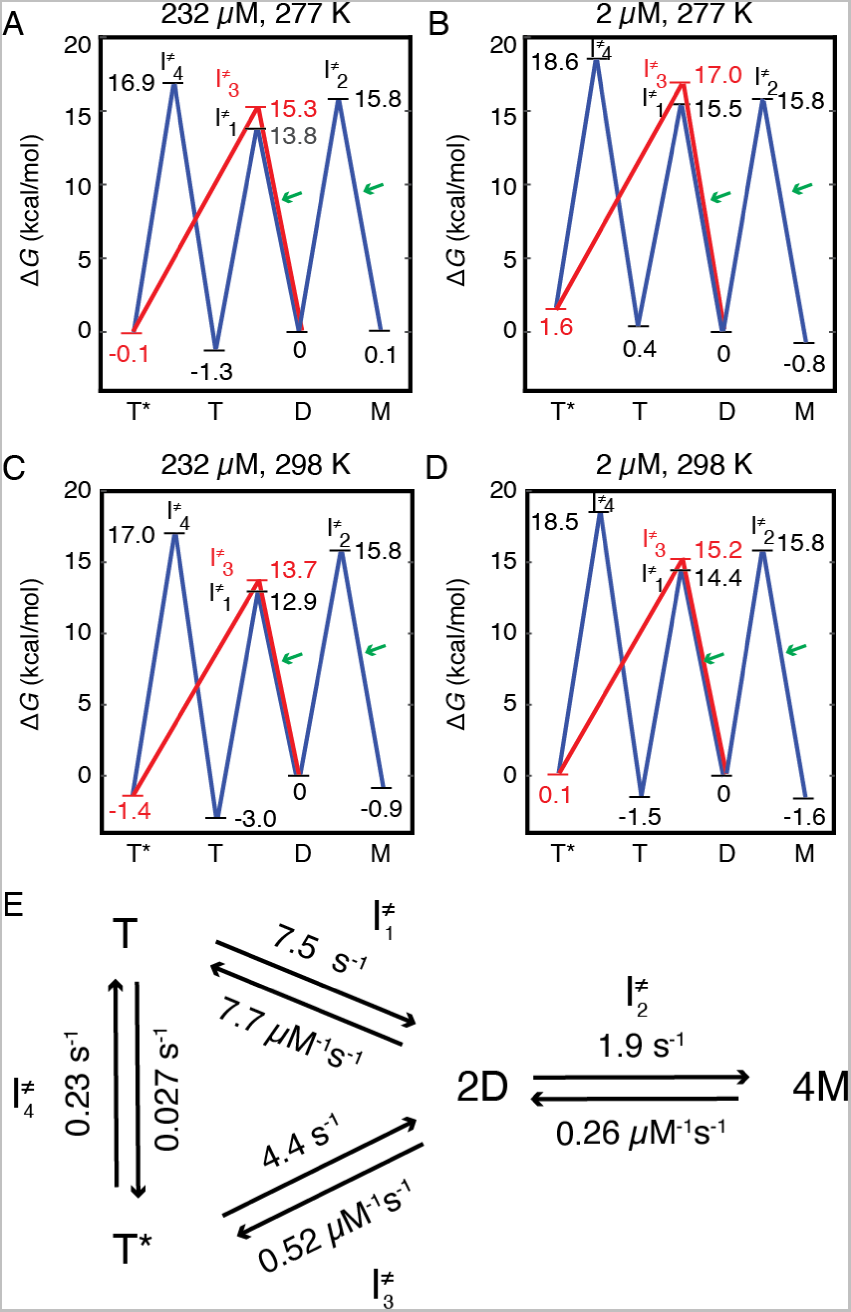
Activation energy diagram for the TDM dissociation pathway of A25T determined from ^19^F-NMR line shape analysis. A–D. Energy diagrams for A25T at 232 µM at 277 K (A) and 298 K (C) and 2 µM at 277 K (B) and 298 K (D). The free energy of the dimer, the common intermediate, is set to 0 kcal/mol as reference. For clarity, the dissociation pathway from T* via 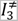 to D is shown in red. The three bi-molecular reactions are labeled with green arrows. E. Schematic depiction of the A25T dissociation pathway. The fitted rate constants at 277 K are labeled with the four intermediates (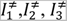 and 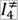). Fitted rate constants at 298 K are shown in Table 2.

**Table 2.** Kinetic parameters fitted by line shape analysis or measured by saturation transfer for the TDM dissociation model of A25T^F^ at 277 and 298 K

| Equilibrium<sup>a</sup> | T ↔ 2D | T* ↔ 2D | D ↔ 2M | T ↔ T* |
| --- | --- | --- | --- | --- |
| Forward rate constant at 277 K | $k_1$<br>$7.5 \pm 0.9 \text{ s}^{-1}$ | $k_5$<br>$4.4 \pm 1.1 \text{ s}^{-1}$ | $k_3$<br>$1.9 \pm 0.4 \text{ s}^{-1}$ | $k_7$<br>$0.027 \pm 0.006 \text{ s}^{-1}$ |
| Reverse rate constant at 277 K | $k_2$<br>$7.7 \pm 1.2 \mu\text{M}^{-1}\text{s}^{-1}$ | $k_6$<br>$0.52 \pm 0.16 \mu\text{M}^{-1}\text{s}^{-1}$ | $k_4$<br>$0.26 \pm 0.05 \mu\text{M}^{-1}\text{s}^{-1}$ | $k_8$<br>$0.23 \pm 0.11 \text{ s}^{-1}$ |
| Forward rate constant at 298 K | $k_1$<br>$13 \pm 1 \text{ s}^{-1}$ | $k_5$<br>$50 \pm 8 \text{ s}^{-1}$ | $k_3$<br>$15 \pm 6 \text{ s}^{-1}$ | $k_7$<br>$0.013 \pm 0.003 \text{ s}^{-1}$ |
| Reverse rate constant at 298 K | $k_2$<br>$\sim 3 \times 10^3 \mu\text{M}^{-1}\text{s}^{-1}$ | $k_6$<br>$\sim 7 \times 10^2 \mu\text{M}^{-1}\text{s}^{-1}$ | $k_4$<br>$1.2 \pm 0.2 \mu\text{M}^{-1}\text{s}^{-1}$ | $k_8$<br>$0.25 \pm 0.07 \text{ s}^{-1}$ |

<sup>a</sup> Rate constants constrained by measured values are in green. Only $k_1$ and $k_5$ are fitting parameters; $k_2$ and $k_6$ are constrained by corresponding $K_d$ values (Table S2).

**Table 3.** Fitted linewidths for the TDM dissociation model of A25T^F^

| <sup>19</sup>F Peak | Line-width (Hz) |  |  |
| --- | --- | --- | --- |
|  | Temperature | Experiment<sup>a</sup> | Modeled<sup>b</sup> |
| T | 277 K | $32 \pm 1$ | $29 \pm 1$ |
| T*/D | 277 K | $32 \pm 2$ | $25 \pm 1$ |
| M | 277 K | $43 \pm 4$ | $38 \pm 1$ |
| T/T*/D | 298 K | $22 \pm 3$ | $19 \pm 1$ |
| M | 298 K | $14 \pm 1$ | $13 \pm 1$ |

<sup>a</sup> From Lorentzian fits of <sup>19</sup>F peaks in spectra recorded at 277 K and concentration below 30 $\mu\text{M}$ , where the T\*/D shoulder and the M peak are significantly populated.
<sup>b</sup> From line-shape analysis based on the TDM model, free of exchange broadening with any other species.

With all rate constants determined for the T*↔2D equilibrium, the MD-calculated ^19^F-NMR line-shape analysis predicts the ratio of diffusion constants of the T* peak over the T peak as 1.11 ± 0.02 at 277 K (see Methods), in excellent agreement with the experimental result of 1.13 ± 0.03 (Figure 2D). This result shows that exchange between T* and D on the second time scale contributes to the measured diffusion constant over a diffusion delay of 0.1 s. The predicted diffusion constant ratio at 298 K (1.02 ± 0.03) also matches the measured ratio of 1.02 ± 0.05, where the dimer species is less populated and thus the diffusion ratio is much closer to 1 since both T and T* are tetrameric. In addition, the kinetic parameters (Table 2) and the linewidth parameters (Table 3) from the line-shape analysis recapitulate ^19^F spectra collected at 1 µM, which is below the concentration range used in the line-shape analysis (Figure S11). Collectively, these comparisons validate the fitting of the ^19^F-NMR line shapes.

Based on the TDM model, the lineshape analysis yields a linewidth for the M peak that is 1.3-fold larger than that of T at 277 K (Table 3), despite M having a lower molecular weight. By contrast, the ratio of the ^19^F *R*_2_ relaxation rate of M over that of T (0.52 ± 0.08), measured with a CPMG pulsing frequency of 4000 s^−1^ for A25T^F^ at 277 K, is comparable to the ratio of linewidths of a F87A^F^ monomer to the tetrameric TTR^F^ (0.60 ± 0.01) or the monomeric F87E^F^ to the tetrameric TTR^F^ (0.63 ± 0.01, Table S4). Thus, the linewidth of the A25T^F^ M species at 277 K in Figure 7A is increased by a process that is quenched by 180° refocusing pulses applied at a high frequency. Exchange between the ground state and an excited state of A25T^F^ M with different ^19^F chemical shifts could lead to such broadening. This was confirmed by ^19^F CPMG dispersion experiments at 277 K, which revealed exchange at a rate of ~780 s^−1^ between the ground and excited state of M. The exchange contributions to relaxation are quenched at a pulsing frequency of 4000 s^−1^ as the fitted ^19^F *R*_2,0_ = 34 ± 3 s^−1^ of A25T^F^ M (Figure S12) is fully consistent with the measured ^19^F *R*_2_ of the A25T^F^ monomer (32 ± 5 s^−1^) as well as the *R*_2_ of the F87A and F87E monomers (Table S4). By comparison, the A25T^F^ T*/D and T peaks do not exhibit relaxation dispersion at 277 K (Figure S12) so that the modeled line width ratio of the T*/D shoulder peak over that of the T peak (0.86 ± 0.04) is within error of the independently measured ^19^F *R*_2_ ratio for A25T^F^ (0.81 ± 0.03, Table S4). The linewidth ratio and ^19^F *R*_2_ ratio are both smaller than 1, indicating a substantial contribution from the smaller dimer species that contributes to the intensity of the T*/D shoulder (Figure 3F). At 298 K, the measured ^19^F *R*_2_ ratio of T*/D shoulder and T peaks of A25T^F^ is 1.0 ± 0.1 (Table S4), consistent with a greatly reduced dimer population at 298 K (Figure 5E) and with the similar translational diffusion coefficients of the T* and T peaks at 298 K (Figure 3D).

The knowledge of all rate constants in the TDM model at two temperatures, 277K and 298K, allows us to construct the free energy landscape for all three dissociation processes (Figure 8). Note that the activation energy for all the unimolecular reactions with first order rate constants *k*_1_, *k*_3_, *k*_5_, *k*_7_ and *k*_8_ are concentration independent. By contrast, the activation free energy for bi-molecular reactions with second order rate constants *k*_2_, *k*_4_ and *k*_6_ (labeled with green arrows in Figure 8A–D) are concentration-dependent. At low concentrations, the activation free energy for these reactions is increased, reflecting reduced oligomerization rates and a more stable monomer relative to other species. Activation enthalpies and entropies determined using transition state theory and the rate constants at two temperatures are listed in Table S5. Of note, the formation of T and T* from D has the most positive activation enthalpy as these two reactions are associated with largest increase in rate constants from 277 to 298 K. The rate constants for conversion from T* to T are nearly the same at both temperatures because the activation enthalpy is close to 0 kcal/mol. Interestingly, the activation enthalpy for the T→T* reaction is negative as the reaction is ca 2-fold faster at 277 than 298 K.

### Molecular dynamics simulations of the A25T dimer

The degeneracy of the ^19^F S85C-BTFA and I84δ1 methyl chemical shifts of D and T* indicate similar environments of the EF loop region in these two species. The F87 side chain is mispacked in the T* species.*^14^* To obtain insights into the structure and dynamics of the A25T dimer we turned to molecular dynamics (MD) simulations. Two types of all-atom simulations were run: isothermal-isobaric ensemble (npt) and canonical ensemble-ensemble (nvt). Multiple independent simulations show that F87 ring flipping occurs on the nanosecond time scale in the A25T dimer (Figures 9A, S14), but is much less frequent in the A25T tetramer (Figure 9B). Corresponding probability contour plots for the dimer and tetramer are shown in S13A, B. Perturbations are observed in the EF loop (Figure 9C) and also in the nearby EF helix and the AB loop in the A25T dimer (Figure S13C, D), which may be related to the loss of stabilizing dimer-dimer interactions that occur across the weak dimer interface in TTR tetramers.

**Figure 9.**
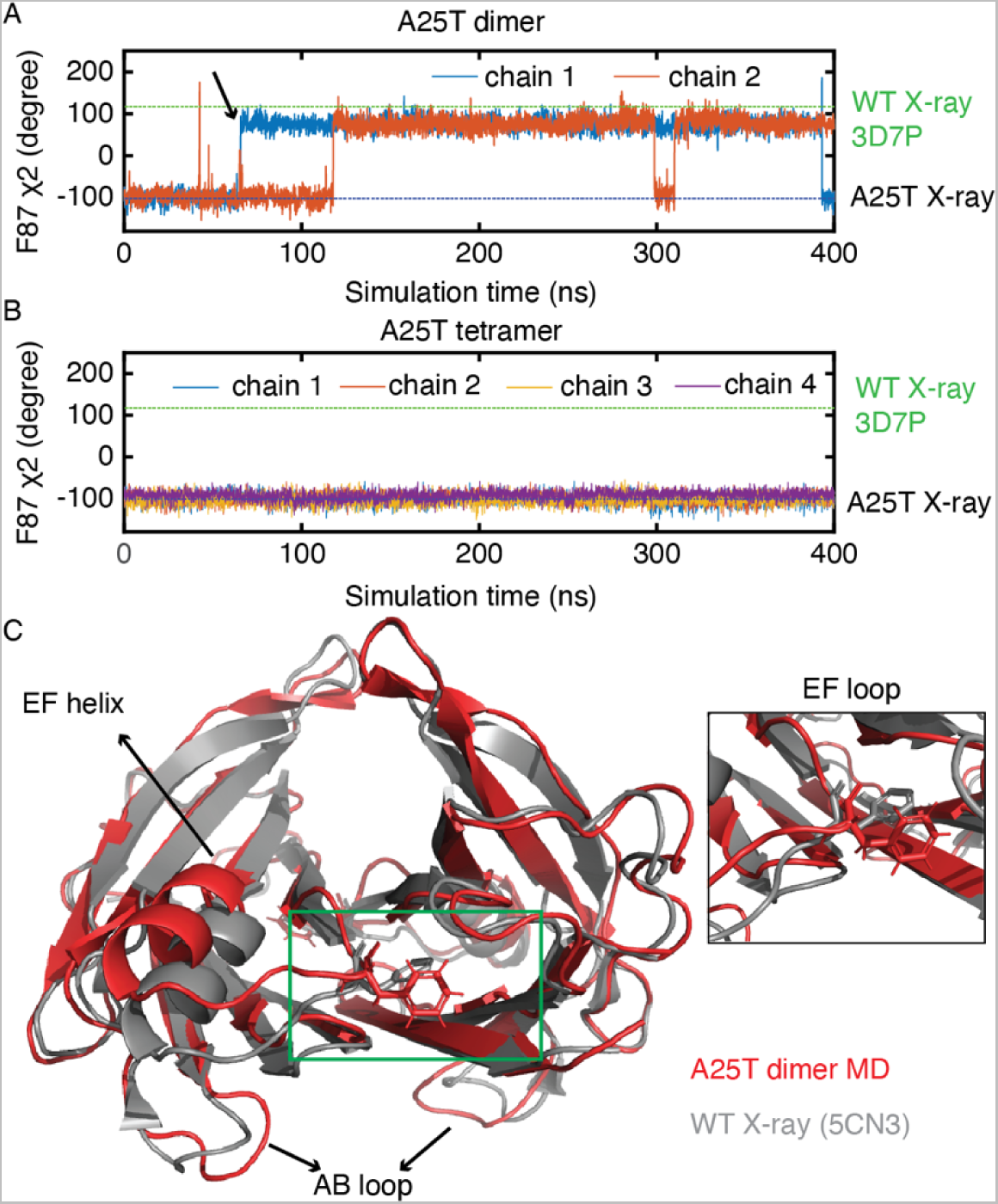
All-atom isothermal-isobaric ensemble (npt) molecular dynamics simulations of the A25T dimer. A–B. Trajectories showing changes in F87 χ2 dihedral angle in A25T dimer (A) and tetramer (B). The F87 χ2 angles in the A25T X-ray structure (black) and a low pH structure of WT TTR (green, PDB 3D7P) are plotted for reference. C. Snapshot from the MD trajectory of the A25T dimer at the time (indicated by an arrow in panel A) when the F87 side chain flips, showing the structural perturbations of the EF helix, EF loop and AB loop.

## Discussion

The dissociation equilibrium of the native TTR tetramer at acidic pH is driven by aggregation of the monomeric aggregation intermediate.*^2, 3, 23^* The quantitative energy landscape of the acid dissociation equilibrium has been measured previously by a ^19^F-NMR aggregation assay.*^17^* At neutral pH, however, the thermodynamic equilibrium constant, the molecular nature of the dissociative intermediate and the kinetics of the dissociation pathway of the TTR tetramer are unknown. Understanding how the TTR tetramer dissociates to less stable monomer at neutral pH is critical due to the neutral physiological pH of blood and cerebrospinal fluid where TTR circulates.

In this work, we used solution NMR based assays to quantitatively map the energetics of the dissociation pathway of a highly destabilized A25T mutant using a tetramer-dimer-monomer (TDM) model (Figure 8A–E). The A25T mutation is located in the AB loop, which forms the weak dimer interface (Figure 1A–B). The enhanced propensity of A25T to dissociate is related to the perturbation propagated to both the weak and strong dimer interfaces; affected residues are either broadened or shifted, as revealed by solution TROSY spectroscopy. Interfacial residues, L17, V20, L110 and V121, a group of residues suggested to play a key role in mediating the dissociation of tetramer to form dimers in MD simulations,*^9^* are broadened in the A25T TROSY spectrum, suggesting enhanced conformational dynamics (Figure 2A–C).

By coupling a BTFA probe at residue S85C, we are able to resolve peaks corresponding to T, T*/D, and M in ^19^F-NMR spectra of the A25T variant under physiological conditions (Figure 3A). The key to our analysis is a global van’t Hoff fit of multiple concentration and temperature titration data sets (Figure 4B–F) and line-shape analysis of the concentration series data sets (Figure 7A–B). The initial intermediate formed on the dissociation pathway is a dimer, the inclusion of which best fits the concentration series of A25T^F^ (Figure 3F and Figure S4) and quantitatively explains the ^19^F-DOSY results (Figure 3D) as it diffuses faster than tetramer and exchanges with the T* under the overlapped shoulder peak on a time scale of seconds (Figure 8E). The degenerate ^19^F chemical shifts of the T* and D peaks suggest that the strong dimer is the dissociative intermediate. The TTR tetramer dissociation pathway via the strong dimer intermediate has been inferred and studied by Cys-crosslinked dimer constructs,*^19, 24^* by surface induced dissociation in gas phase by native mass spectrometry*^25^* and by MD-based free energy calculations.*^9, 10^* Our results of dimer-mediated dissociation are obtained in neutral pH solution under equilibrium conditions and are consistent with these previously reported results. Numerical simulations have suggested that the dimer-mediated formation of tetramer (dimer of dimers) can reduce amounts of protein trapped as assembly intermediates and is employed by all homo-tetrameric proteins surveyed.*^26^* In this regard, the homotetrameric TTR is no exception.

The equilibrium population of the dimer is less than 5% at 298 K when the A25T concentration is above 0.1 µM (Figure 5E). This small population highlights the need of combining a highly sensitive ^19^F trifluoromethyl probe and cryoprobes to study the dimeric species experimentally. The similar environments experienced by ^19^F and I84δ1 probes in D and T* offer structural insights into the otherwise elusive dimer species. As suggested by MD simulations, the F87 side ring flips more frequently on the nanosecond time scale in the A25T dimer than in the tetramer (Figure 9A–B). These ring flipping events are likely coupled with altered EF loop conformations that are common in T*. Thus the EF loop conformations are perturbed in D and T* compared to that in the native tetramer in which the F87 ring is correctly packed,*^14^* resulting in a slightly upfield shifted ^19^F chemical shift in the shoulder peak from the main T peak (Figure 3A–C). The mispacked T* state is observed under physiological conditions (Figure 3A) and all the species can be converted to T with tafamidis within 1 hour (Figure 3B), consistent with all the interconversion rate constants on the second to minute time scale (Figure 8E). Interestingly, the energy difference between the T and T* states of A25T^F^ at 298 K is 1.6 kcal/mol (Figure 8C–D), which is identical to that of the parental mutant TTR^F^ (1.6 kcal/mol or a T* population of ~7%, as previously described*^14^*). This similarity indicates that the A25T mutation does not change the energetic difference between the two forms of the TTR tetramer.

The quantitative insights from our analysis allow us to place our results in the context of existing TTR literature. First, the Δ*H* and Δ*S* of the forward T↔4M equilibrium of A25T are more negative than reported values for two EF helix mutants, K80E and K80D.*^5^* However, Δ*G* at 298 K is similar for all three destabilized mutants, ~7–8 kcal/mol less than for WT TTR (Table S6). Second, the rate constants for the D↔2M transition have been reported for WT TTR refolded upon dilution from denatured monomers as 9–588 s^−^*^1^* (*k*_3_) and 0.3–20 µM^−1^s^−1^ for (*k*_4_) at 298 K*^27^*. Our rate constants *k*_3_ = 15 s^−1^ and *k*_4_ = 1.2 µM^−1^s^−1^ measured for A25T^F^ at 298 K fall within these ranges. Third, the T* unfolding rate constant is 0.26 h^−1^ and that for T is 0.066 h^−1^ in TTR^F^ at 298 K*^14^*. Because urea cannot denature the TTR tetramer*^19^*, the urea unfolding rate is thus associated with dissociation of tetramer. For A25T^F^ at 298 K, the rates of dimer formation by dissociation of T* and T (*k*_5_ and *k*_1_) are 50 and 13 s^−1^, respectively, much faster than the corresponding urea unfolding rate constants for T and T* in TTR^F^. This is consistent with the A25T mutant as a highly kinetically destabilized variant.*^4, 11^* Interestingly, the *k*_5_/*k*_1_ ratio for T* and T A25T^F^ is 3.8 ± 0.7, closely mirroring the 3.9-fold increase of urea unfolding rate constants of T* and T in TTR^F^. This similarity suggests that the A25T mutation likely affects the dissociation processes of T and T* similarly.

Based on subunit exchange assays at equilibrium,*^28^* WT TTR dissociates faster at 277 K (0.089 h^−1^)*^20^* than 298 K (~0.037 h^−1^), a 2.4-fold increase at the lower temperature. For A25T, only *k*_7_ (T → T*) shows an increase at 277 compared to 298 K (Table 2), which, interestingly, also shows a 2.1 (± 0.7)-fold increase. This anti-Arrhenius behavior is related to the negative 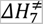 (Table S5). The *k*_7_ = 0.013 ± 0.003 s^−1^ (0.8 ± 0.2 min^−1^) is on the same time scale as the dissociation rate constant of A25T at 298 K as 0.3 min^−1^, which was extrapolated from urea denaturation experiments to [urea] = 0 M.*^11^* Without urea, the apparent overall dissociation rate constant is comparable to the subunit exchange rate.*^29^* These comparisons suggest the slowest first order rate constant *k*_7_ plays a role in affecting subunit exchange rates of TTR as well as its temperature dependence at neutral pH.

In summary, we demonstrate an integrated method to allow simultaneous determination of all thermodynamic and kinetic parameters of the TDM model for A25T dissociation at neutral pH. Our approach reveals free energies as well as activation barriers for all equilibrium steps, including ones involving lowly populated species. We expect this set of tools to be applicable to a wide range of protein systems of interest.

## Supporting information

The supporting information contains materials and methods, supplementary tables 1-6 and supplementary figures 1-14.

## Acknowledgements

We thank Gerard Kroon and Maria Martinez-Yamout for expert assistance in NMR experiments and molecular biology, respectively, and Euvel Manlapaz for technical support.

