## Supplementary Material for "Probing the dissociation pathway of a kinetically labile transthyretin mutant"

### Materials and Methods

#### Protein expression

The expression, purification and $^{19}$F labeling of non-tagged A25T$^{F}$ was performed as previously described[1]. Non-tagged A25T used in X-ray crystallography and I73V, I84V and F87E-V122I used in methyl assignment were purified as previously described[2]. Other A25T plasmids were based on Histag-tevG-A25T in pET21a plasmid and expressed in a BL21(DE3)-DNAY strain. Cells were induced with 1 mM IPTG at 310 K for 4 hours when the OD reached 0.8–1.0. An additional Gly was introduced after the tev protease cleavage site (ENLYFQG) to facilitate the tev protease cleavage. To purify TTR constructs with the Histag, the lysed protein was loaded on 5 mL Ni cOmplete resin (Roche) equilibrated in 20 mM Tris/HCl and 1 M NaCl at pH 8.0. The TTR constructs were eluted in 15-25 mL 20 mM Tris/HCl, 1 M NaCl and 200 mM imidazole at pH 8.0 after washing with 250 mL 20 mM Tris/HCl, 1 M NaCl and 5 mM imidazole at pH 8.0. The tev protease digestion was carried with a 1:100 tev:TTR molar ratio in a dialysis membrane (6–8 k Da molecular weight cutoff) against the Ni resin elution buffer (20 mM Tris/HCl, 1M NaCl, 200 mM imidazole at pH 8.0) for 1 day at 298 K or 20 mM Tris/HCl, 100 mM NaCl and 3 mM DTT at pH 8.0 overnight at 298 K. The reaction mixture was then reloaded to the Ni resin, eluted in 20 mM Tris/HCl and 1 M NaCl at pH 8.0, buffer exchanged to NMR buffer (10 mM phosphate potassium, 100 mM KCl at pH 7.0) and frozen at -80 $^{o}$C before use.

#### SDS PAGE analysis

Samples of 30 µM TTR mutants were mixed with 6× sodium dodecyl sulfate (SDS) loading buffer with DTT. Prior to loading to a Mini-PROTEAN TGX Precast Polyacrylamide gel (Bio-Rad), samples were boiled for 90 s. Unboiled samples were loaded as controls and the gel was stained by Coomassie blue.

#### Size exclusion column chromatography

A prepacked Superdex 75 column (GE Lifescience, 10 × 285) was equilibrated in NMR buffer and run at 0.5 mL/min at 298 K using AKTA Pure. A solution of 100 µL 4 µM A25T$^{F}$ filtered by 0.22-µm filter was injected. The dilution factor was calculated as a ratio between the M peak width (1 or 2 s.d. from a Gaussian fit of the M absorbance peak) and the injected volume (100 µL), leading to 0.2 or 0.1 µM in-column concentration.

#### $^{1}$H, $^{13}$C-HMQC and $^{1}$H, $^{15}$N-TROSY spectroscopy

Standard $^{1}$H,$^{13}$C-HMQC spectra were acquired using a Bruker Avance 600 spectrometer with 2k and 128 complex points in the $^{1}$H and $^{13}$C dimensions, respectively, and spectral widths of 16 and 36 ppm. The carrier offset was set at 4.7 and 20 ppm for the $^{1}$H, $^{13}$C dimensions, respectively. Samples were uniformly $^{13}$C-labeled. Standard $^{1}$H,$^{15}$N-TROSY spectra were acquired using a Bruker Avance 800 spectrometer with 2k, and 256 complex points in the $^{1}$H and $^{15}$N dimensions and spectral widths of 16 and 32 ppm. The carrier offset was set at 4.7 and 118 ppm for the $^{1}$H, $^{15}$N dimensions, respectively. A recycle delay of 1 s was used for both HMQC and TROSY experiments. For backbone assignments, deuterated A25T at 0.4 mM was used in the NMR buffer with 2 mM TCEP, and amide or Cα peaks that are not broadened were assigned using standard trHNCA/trHN(CO)CA spectra. Ile δ1 methyls in the A25T tetramer were assigned using I26V, I68V, I73V and I84V mutations. Of note, A25T-I107V and A25T-I73V are insoluble; the latter construct is consistent with a previous report that I73 is important for stabilization of the tetramer[3]. The Ile δ1 methyls in the A25T monomer were assigned by transferring Ile δ1 assignments from the spectrum of a monomeric V122I-F87E whose Ile δ1 side chains were assigned by using H(CCO)NH, (H)C(CO)NH, H(C)CH-COSY, (H)CCH-COSY, H(C)CH-TOCSY and (H)CCH-TOCSY. Peak intensity and volume were measured using Sparky[4]. Random coil Cα chemical shifts were calculated by POTENCI[5].

**$^{19}$F-NMR data acquisition and processing**

The 1D $^{19}$F-NMR spectra were collected using Bruker Avance 600 or Avance 700 spectrometers as previously described[1]. A 1-Hz exponential line-broadening factor was applied to the free induction decay, which was then zero-filled to 16k before Fourier transformation. Temperature titrations (at 277, 280, 283, 286, 289, 292, 295 and 298 K) were performed for 30 μM A25T$^F$ using the Avance 600 spectrometer or for 6 and 230 μM A25T$^F$ using theAvance 700 spectrometer. The concentration dilution of A25T$^F$ monitored by $^{19}$F-NMR was carried using the Avance 700 spectrometer at 277 and 298 K. A25T$^F$ at 30 μM was incubated with 50 μM tafamidis at 277 K for 1 hour prior to the $^{19}$F-NMR measurement at 281 K using the Avance700 spectrometer.

**$^{19}$F-NMR relaxation experiments**

$^{19}$F-NMR $R_1$ and $R_2$ relaxation rates (Table S3 and S4) were measured as previously described in Ref. 2 and Ref. 6 respectively for A25T$^F$ using the Avance 600 spectrometer.

**$^{19}$F-NMR diffusion ordered spectroscopy**

$^{19}$F-NMR DOSY experiments were performed as previously described[1]. Typically, ten experiments with ten evenly spaced relative z-gradient strengths from 5% to 50% were collected overnight.

**The TDM model for A25T dissociation**

In this model that describes the dissociative pathway of the A25T tetramer via a dimer to form monomers (TDM), three equilibrium constants ( $K_\mathrm{d}$ ) are defined as follows:

$$K_{\mathrm{d}1} = \frac{[D]^2}{[T]} \qquad \text{Eq. 1}$$

$$K_{\mathrm{d}2} = \frac{[M]^2}{[D]} \qquad \text{Eq. 2}$$

$$K_{\mathrm{d}3} = \frac{[D]^2}{[T^*]} \qquad \text{Eq. 3}$$

where T, T*, D and M stand for the native tetramer, mispacked tetramer, dimer and monomer, respectively (Scheme 1). The concentrations in Eq. 1 to 3 are molar concentrations of each species, not protomer concentrations. The $K_{\mathrm{d}4}$ between [T] and [T*] can be calculated as $K_{\mathrm{d}4} = [T]/[T^*] = K_{\mathrm{d}3}/K_{\mathrm{d}1}$.

By mass conversion, we have:

$$4[T] + 4[T^*] + 2[D] + [M] = c_t \qquad \text{Eq. 4}$$

where $c_\mathrm{t}$ is the total concentration of TTR protomer quantified by an extinction coefficient of 18,450 $M^{-1}$ $cm^{-1}$ at 280 nm. By rearranging this mass conversion equation, the following quartic polynomial is derived:

$$\left(\frac{4}{K_{\mathrm{d}1}K_{\mathrm{d}2}^2} + \frac{4}{K_{\mathrm{d}3}K_{\mathrm{d}2}^2}\right) f_\mathrm{M}{}^4 + \frac{2}{c_t^2 K_{\mathrm{d}2}} f_\mathrm{M}{}^2 + \frac{1}{c_t^3} f_\mathrm{M} - \frac{1}{c_t^3} = 0 \qquad \text{Eq. 5}$$

where the monomer population fraction $f_\mathrm{M}$ is defined as $[M]/c_\mathrm{t}$, equivalent to the $^{19}$F peak area of monomer relative to the total $^{19}$F peak area.

A series of A25T$^F$ samples at different concentrations were made at 298 K prior to measurements at 277 K and 700 MHz in NMR buffer. There are three resolved $^{19}$F peaks in spectra of A25T$^F$ recorded over a range of concentrations. These were assigned as the correctly packed tetramer (T), a shoulder peak and monomer M (see Figure 3E). At each concentration, the $^{19}$F spectrum was fitted by three Lorentzian functions to extract the relative population of each species. In the TDM model, the $^{19}$F peak position of the dimer species was assumed to overlap with that of the mispacked tetramer, T*, under the same shoulder T* peak. The solution of $f_M$ as a function of $c_t$ was numerically solved by the *roots* function in MATLAB for a set of floating $K_d$ values. Then the relative populations of T, T* and D were computed based on Eqs. 1–5, and the residuals compared to experimental populations, determined from the Lorentzian fits, were iteratively minimized as previously described in Ref. 7. Unless otherwise noted, the *fminsearch* function in MATLAB was used in minimizing the fitting residues between model and data, and the fitting uncertainty was reported as one standard deviation from 50 bootstrapped datasets.

**van't Hoff analysis of the TDM dissociation pathway of A25T$^F$**

The temperature dependence of $K_{d1}$, $K_{d2}$, and $K_{d3}$ is related to $\Delta H_{1,2,3}$ and $\Delta S_{1,2,3}$ by the van't Hoff equation:

$$\ln K_i = -\frac{\Delta H_i}{RT} + \frac{\Delta S_i}{R} \qquad \text{Eq. 6}$$

where $R$ is the ideal gas constant and $T$ is the temperature for the equilibria defined in Eq. 1 to 3 for $i$ = 1, 2 or 3. A global fit for measured A25T populations (T, T*, D and M) as a function of temperature and concentration was performed to fit $\Delta H_{1,2,3}$ and $\Delta S_{1,2,3}$.

**$^{19}$F-NMR Line shape analysis**

The $^{19}$F-NMR line shape analysis was based on numerically solving the Bloch-McConnell equation for describing chemical exchange in an isolated spin system without scalar coupling[8] as:

$$\frac{d\mathbf{N}(t)}{dt} = (i\mathbf{L} - \mathbf{R} + \mathbf{K})\mathbf{N}(t) \qquad \text{Eq. 7}$$

where the transverse magnetization evolution vector $\mathbf{N}(t)$ is defined as $[N_T(t);\ N_{T^*}(t); N_D(t); N_M(t)]$. The elements of the diagonal Liouvillian matrix $\mathbf{L}$ are given by $\delta_{ij}\omega_{ij}$ where $\omega$ stands for the chemical shift of the four A25T species in the same order defined in $\mathbf{N}(t)$. Similarly, the elements in the diagonal relaxation matrix $\mathbf{K}$ are $\delta_{ij}R_{2,ij}$ where $R_2$ is the transverse relaxation constant for each species in the absence of exchange with other species. The kinetic exchange matrix $\mathbf{K}$ for the TDM dissociation model is:

$$\mathbf{K} = \begin{pmatrix} -k_1 - k_7 & k_8 & 2k_2[D] & 0 \\ k_7 & -k_5 - k_8 & 2k_6[D] & 0 \\ k_1 & k_5 & -2k_2[D] - 2k_6[D] - k_3 & 2k_4[M] \\ 0 & 0 & k_3 & -2k_4[M] \end{pmatrix} \qquad \text{Eq. 8}$$

where the rate constants are defined in Scheme 1.

The following constraints were used in solving Eq. 7: $K_{d1} = k_1/2k_2$, $K_{d2} = k_3/2k_4$ $K_{d3} = k_5/2k_6$ and $K_{d4} = [T]/[T^*] = K_{d3}/K_{d1}$. The chemical shift $\omega$ in the matrix $\mathbf{L}$ is fixed at invariant peak positions of T, T*/D and M extracted by 3-state Lorentzian fits across all concentrations (Figure S9). Furthermore, $k_3/k_4$ at an intermediate concentration of 30 μM A25T$^F$ was measured at 277 K by $^{19}$F saturation transfer (Figure S10). At 298 K, $k_3/k_4$ was measured at a low concentration (6 μM) to favor the M species and $k_7/k_8$ was measured at a high concentration (260 μM) to suppress the D population under the shoulder T* peak. These measured rate constant constraints are colored in green in Table 2. Due to a higher D population under the shoulder T* peak at 277 K, $k_7/k_8$ at 277 K was estimated based on measurements at 298 K and transition state theory (see following two sections below). Therefore, only two rate constants ($k_1$ and $k_5$) were floated at both temperatures. Floating all four linewidth yields unconstrained results, indicating overfitting. To constrain the fits with a minimal number of floating parameters, three linewidths that are free of inter-species exchange were floated for the three observed $^{19}$F peaks for T, T*/D and M at 277 K. At 298 K, two linewidths were floated for T/T*/D and M respectively due to similar $^{19}$F $R_2$ values for the T and T*/D peak measured at 600 MHz (Table S4). In summary, there are five and four floating parameters at 277 and 298 K, respectively. The eigen method from Ref. 9 was used to solve the Bloch-McConnell equation and the real parts of $\mathbf{N}(t)$ were Fourier transformed and compared to experimental line shapes. The fitting errors were normalized using a bounded variant of the *fminsearch* function in MATLAB. The fitted $k_1$ and $k_5$ are listed in Table 2 and fitted line widths are in Table 3.

### $^{19}$F-NMR saturation transfer experiments

$^{19}$F saturation transfer experiments were performed at 277 K and 600 MHz using 30 μM A25T$^{F}$. The T*/D peak was saturated using a weak 60 dB pulse over a range of mixing times from 25 ms to 2 s. The recycle delay was set at 4 s. The $^{19}$F carrier was placed at -84.833 and -85.137 ppm for on and off resonance experiments, respectively. This way, the spectral separation from the M peak to the two carrier positions was identical. The peak area of M was used in quantification, normalized by a control experiment without the saturation mixing period. A total of 512 scans were collected for 10 1D spectra with varying mixing times for both on and off conditions. All experiments were collected in an interleaved manner to minimize time-dependent sample changes. The data were analyzed per Ref. 10 where the rate of dimerization of M ($k_{\mathrm{MD}} = 2k_4[M]$) is calculated as:

$$k_{\mathrm{MD}} = \frac{1}{T_1^{\mathrm{M}}}\left(\frac{I_{t=0}^{\mathrm{M}}}{I_{t=\infty}^{\mathrm{M}}} - 1\right) \qquad \text{Eq. 9}$$

where $T_1^{\mathrm{M}}$ is the longitude relaxation time of M, $I_{t=0}^{\mathrm{M}}$ is the intensity of the M peak without mixing and $I_{t=\infty}^{\mathrm{M}}$ is the steady-state intensity of the M peak with an infinite mixing period. In practice for the A25T system, a 2-s mixing time was found to be sufficient (Figure S10). At a total concentration of 30 μM, [M] = 2.5 μM based on van't Hoff analysis.

Saturation transfer experiments were also performed at 298 K, using a low concentration (6 μM) of A25T$^{F}$ to enhance the relative M population. The $^{19}$F carrier was placed at -84.316 and -84.634 ppm for on and off resonance experiments, respectively. A total of 4000 scans were collected for 10 transfer transients for both on and off conditions at 700 MHz.

A saturation transfer experiment for 260 μM A25T$^{F}$ at 298 K was performed at 600 MHz to measure the T↔T* exchange. The T peak was saturated and the T* peak was probed. The frequency offset of the $^{19}$F carrier was set at -84.148 and -84.398 ppm for the on and off resonance experiments, respectively. A total of 280 scans were collected. This experiment measures the rate constant for T* to convert to T ($k_8$):

$$k_8 = \frac{1}{T_1^{\mathrm{T}^*}}\left(\frac{I_{t=0}^{\mathrm{T}^*}}{I_{t=\infty}^{\mathrm{T}^*}} - 1\right) \qquad \text{Eq. 10}$$

The reverse rate constant from T to T* ($k_7$) is too slow to be accurately measured by $^{19}$F saturation transfer experiments and was determined from the detailed kinetic balance between T and T*:

$$k_7 = \frac{[T^*]}{[T]} k_8 \qquad \text{Eq. 11}$$

All measured results are colored in green and listed in Table 2.

**Transition state theory analysis**

The activation free energy $\Delta G_i^{\neq}$ of the T↔T* exchange based on transition state theory is:

$$\Delta G_i^{\neq} = RT[\ln(k_B T/h) - \ln(k_i)] \qquad \text{Eq. 12}$$

where $k_i$ is the measured rate for $i$ = 7 or 8 (see Scheme 1), $k_B$ is the Boltzmann constant, and $h$ is the Planck constant.

The activation enthalpy $\Delta H_i^{\neq}$ and entropy $\Delta S_i^{\neq}$ for steps $i$ = 7 or 8 are constrained by the following equations due to the closed thermodynamic cycle among T, T* and the transition state:

$$\Delta H_7^{\neq} = \Delta H_8^{\neq} + \Delta H_1 - \Delta H_3 \qquad \text{Eq. 13}$$

$$\Delta S_7^{\neq} = \Delta S_8^{\neq} + \Delta S_1 - \Delta S_3 \qquad \text{Eq. 14}$$

where $\Delta H_i$ and $\Delta S_i$ with $i$ = 1 or 3 are derived from the van't Hoff analysis (Table 1).

$\Delta H_i^{\neq}$ and $\Delta S_i^{\neq}$ can be related to $\Delta G_i^{\neq}$ by

$$\Delta G_i^{\neq} = \Delta H_i^{\neq} - T\Delta S_i^{\neq} \qquad \text{Eq. 15}$$

Based on measured values of $k_7$ and $k_8$ at 298 K, $\Delta H_i^{\neq}$ and $\Delta S_i^{\neq}$ for $i$ = 7 or 8 can be numerically solved by the *fsolve* function in MATLAB given Eqs. 12–15. This allows determination of $k_7$ and $k_8$ at 277 K, where overlap between the D and T* peaks and the broadness of the M peak complicate the direct measurement of $k_7$ and $k_8$.

The activation enthalpy $\Delta H_i^{\neq}$ and entropy $\Delta S_i^{\neq}$ for $i$ = 1, 3 and 5 are determined by $k_i$ from the line-shape analysis at 277 K and 298 K (Table 2) and the corresponding reverse $\Delta H_i^{\neq}$ and entropy $\Delta S_i^{\neq}$ for $i$ = 2, 4 and 6 are calculated as follows thanks to closed thermodynamic cycles:

$$\Delta H_i^{\neq} = \Delta H_{i+1}^{\neq} + \Delta H_i \qquad \text{Eq. 16}$$

$$\Delta S_i^{\neq} = \Delta S_{i+1}^{\neq} + \Delta S_i \qquad \text{Eq. 17}$$

where $\Delta H_i$ and $\Delta S_i$ for $i$ = 1, 2 or 3 are from the van't Hoff analysis (Table 1). The $\Delta H_i^{\neq}$ and $\Delta S_i^{\neq}$ values are summarized in Table S5.

**$^{19}$F $R_2$ relaxation dispersion analysis**

Constant-time $^{19}$F CPMG relaxation dispersion[11] data were acquired for 30 μM A25T$^F$ at 277 K using the Avance 600 spectrometer. The constant time was 10 ms. The effective $^{19}$F $R_2$ relaxation rate constant ($R_2^{\text{eff}}$) of M was fitted using the fast exchange Luz-Meiboom equation[12]:

$$R_2^{\text{eff}} = R_2^0 + \frac{\Theta}{k_{\text{ex}}}\left[1 - \frac{2}{k_{\text{ex}}\tau}\tanh\left(\frac{k_{\text{ex}}\tau}{2}\right)\right] \qquad \text{Eq. 18}$$

where $\Theta = p_A p_A \Delta\omega^2$, $R_2^0$ is the exchange-free transverse relaxation rate constant, $k_{\text{ex}}$ is the exchange rate constant between the ground state and the excited state of M and $\tau$ is the delay between 180° pulses in the CPMG pulse train.

**$^{19}$F-NMR DOSY simulations**

The DOSY simulation for fast exchange between T* and D was performed according to Ref. 13.

Briefly, the magnetization of the $^{19}$F shoulder T* peak ($N(t,g)$) is the sum of the magnetization from the T* and D species that undergo exchange with rate constants defined as $k_5$ and $2k_6[D]$ (see Scheme 1). The time- and z-gradient dependent $N(t,g)$ is:

$$N(t,g) = \left[\frac{N_0}{2} + \frac{\Lambda}{2\Delta}\right] e^{[(-\sigma+\Delta)t]} + \left[\frac{N_0}{2} - \frac{\Lambda}{2\Delta}\right] e^{[(-\sigma-\Delta)t]} \qquad \text{Eq. 19}$$

where the variable $t$ is the diffusion delay (100 ms in the $^{19}$F DOSY experiments), and $g$ is the relative z-gradient. Other parameters are defined as follows:

$$\Lambda = \emptyset(N_{\mathrm{D},0} - N_{\mathrm{T}^*,0}) + N_{\mathrm{T}^*,0}k_5 + 2k_6[D]N_{\mathrm{T}^*,0}$$

$$\sigma = \frac{1}{2}(k_5 + 2k_6[D] + D_{\mathrm{T}^*}g^2 + D_{\mathrm{D}}g^2)$$

$$\emptyset = \frac{1}{2}(k_5 - 2k_6[D] + D_{\mathrm{T}^*}g^2 - D_{\mathrm{D}}g^2)$$

$$\Delta = \sqrt{\emptyset^2 + 2k_5k_6[D]}$$

$$N_0 = N_{\mathrm{T}^*,0} + N_{\mathrm{D},0}$$

where $D_{\mathrm{T}^*}$, the translational diffusion coefficient of the T* species, is set as 1 (unitless) for reference and $D_{\mathrm{D}}$ = 1.24 calculated as a ratio of the diffusion coefficient of dimer and tetramer estimated using the Stoke-Einstein equation as previously described by Ref. 1. $N_{\mathrm{T}^*,0}$ and $N_{\mathrm{D},0}$ are equilibrium magnetization derived from the TDM model at either 298 or 277 K at matching concentrations, $k_5$ and $k_6$ were constrained using the fitted results from the line shape analysis at either 298 or 277 K (Table 1). The first ten points of simulated relative gradient strengths were used in linear fits where the logarithm of the intensity was fitted against $g^2$ to extract slopes as simulated diffusion constant ratios to compare with experimental ratios.

**X-ray crystallography and structural modeling**

The A25T variant was prepared for crystallization using the published method that uses ammonium sulfate.[14] Crystallization screens were set up using the sitting drop vapor diffusion method. A25T at 10 mg/mL was crystalized in 0.1 M sodium cacodylate (pH 6.5), 0.2 M calcium acetate and 40 % (w/v) PEG 300 at 277 K. Diffraction data were processed using HKL-2000[15]. Molecular replacement was performed using Phaser[16] with TTR structure PDB:2ROX as a search model. Models were refined with phenix.refine[17], refmac[18], and Coot[19]. Data collection and refinement statistics are listed in Table S1.

**All-atom molecular dynamics simulations**

Our A25T X-ray structure was used as a starting dimer model for molecular dynamics (MD) simulations using the AMBER ff14SB[20] force field and AMBER 16[21] software. The symmetry imposed by X-ray crystallography was used to create a tetrameric A25T as an initial tetramer conformation using PyMOL (2.5.0). The protonation states of four histidine residues at pH 7.0 were determined using the H+++ server[22] and our A25T X-ray structure. The preparation of the X-ray structures was performed as previously described[2]. Briefly, each structure was embedded in a cuboid TIP3 water[23] box larger than TTR by at least 10 Å from the box edges. Periodic boundary conditions were then applied. The net charge of the solvated system at pH 7.0 was neutralized by $K^+$ and the ionic strength was set to 100 mM as in the NMR buffer using $K^+$ and $Cl^-$. The solvated systems were then heated from 0 to 298 K during 40 ps MD simulations with a time step of 2 fs using a Langevin thermostat with 1 ps$^{-1}$ collision frequency, during which a weak harmonic restraint of 10 kcal/mol/Å$^2$ was applied on all protein heavy atoms. Next, for equilibration, two sequential isothermal-isobaric ensemble (npt) runs of 2 ns each were performed at 298 K and 1 atm with 10 and 1 kcal/mol/$Å^2$ restraints on protein heavy atoms, respectively. The equilibration runs were performed using a Langevin thermostat with 1 $ps^{-1}$ collision frequency with isotropic position scaling and a pressure relaxation time of 1 ps. A final equilibrium npt ensemble simulation of 1 ns at 298 K and 1 atm was performed without any force restraints on protein heavy atoms. All bond lengths involving hydrogen atom were constrained at equilibrium lengths by the SHAKE algorithm[24]. The cutoff radius for van der Waals and real-space particle mesh Ewald electrostatics was set at 10 Å. Productive MD simulations of 400 ns were carried out starting with the npt ensembles at 298 K. MD snapshots were saved every 0.1 ns for analysis using CPPTRAJ[25] and MATLAB. The canonical ensemble ensemble simulations at 298 K using a Berendsen thermostat with a coupling time of 10 ps were also performed (Figure S14).

**Table S1** X-ray data collection and refinement statistics for A25T

| **Mutant** | **A25T** |
|---|---|
| **Data Collection** | |
| **Beamline** | ALS 5.0.3 |
| **Wavelength (Å)** | 0.9765 |
| **Resolution (Å)** | 39.0-1.63 (1.66-1.63) |
| **Space Group** | P $2_1$ $2_1$ 2 |
| **Unit Cell (Å)** | a=43.78 b=86.03 c=64.65 |
| **Total reflections[a]** | 241,435 (10,422) |
| **Unique reflections** | 30,727 (1455) |
| **Redundancy** | 7.9 (6.9) |
| **Completeness (%)** | 98.8 (95.5) |
| **<I>/<σI>** | 22.2 (1.1) |
| **$R_{merge}$ (%)[b]** | 10.7 (177) |
| **$R_{meas}$ (%)[c]** | 11.5 (191) |
| **$R_{pim}$ (%)[d]** | 4.0 (70.5) |
| **$CC_{1/2}$ (%)[e]** | 87.1 (41.5) |
| **Refinement Statistics** | |
| **Resolution (Å)** | 39.0-1.63 |

| | |
|---|---|
| | (1.68-1.63) |
| **$R_{work}$/$R_{free}$ (Å)** | 19.6/23.6 |
| **Number of reflections in refinement (work/free)** | 29,134/1544 |
| **Number of non-H protein atoms** | 1852 |
| **Number of water molecules** | 201 |
| **Number of protein residues** | 249 |
| **RMS (bonds)** | 0.008 |
| **RMS (angles)** | 1.04 |
| **Ramachandran: favored, outliers (%)** | 97.4, 0 |
| **Clashscore[f]** | 6.3 |
| **Wilson B (Å$^2$)** | 18 |
| **Average B (Å$^2$)** | 26 |
| **Protein** | 25 |
| **Chain A** | 24 |
| **Chain B** | 26 |
| **Water** | 34 |

[a] Numbers in parentheses are for highest resolution shell
[b] $R_{merge} = \Sigma_{hkl} \Sigma_{i=1,n}|I_i(hkl) - \langle I(hkl)\rangle| / \Sigma_{hkl} \Sigma_{i=1,n} I_i(hkl)$
[c] $R_{meas} = \Sigma_{hkl} \sqrt{(n/n-1)}\, \Sigma_{i=1,n}|I_i(hkl) - \langle I(hkl)\rangle| / \Sigma hkl\, \Sigma_{i=1,n} I_i(hkl)$
[d] $R_{pim} = \Sigma_{hkl} \sqrt{(1/n-1)}\, \Sigma_{i=1,n}|Ii(hkl) - \langle I(hkl)\rangle| / \Sigma hkl\, \Sigma_{i=1,n} I_i(hkl)$
[e] $CC_{1/2}$ = Pearson Correlation Coefficient between two random half datasets
[f] Number of unfavorable all-atom steric overlaps ≥ 0.4. per 1000 atoms

**Table S2** Statistically similar $K_d$ of A25T$^F$ at 277 K by fitting different datasets using the van't Hoff equation.

| Data sets | $K_{d1}$ (μM) [a] | $K_{d2}$ (μM) [b] | $K_{d3}$ (μM) [c] |
|---|---|---|---|
| One concentration series at 277 K (A) | 0.5 ± 0.1 | 4.0 ± 0.3 | 3.3 ± 0.6 |
| Three temperature series at 6, 30 and 260 μM separately (B) | 0.5 ± 0.1 | 3.7 ± 0.5 | 4.0 ± 1.1 |
| (A) + (B) | 0.5 ± 0.1 | 3.9 ± 0.2 | 3.7 ± 0.5 |
| (A) + (B) + one more concentration series at 298 K[d] | 0.5 ± 0.1 [2 ± 1 nM] | 3.7 ± 0.3 [6 ± 2 nM] | 4.1 ± 0.7 [34 ± 21 nM] |

[a] $K_{d1} = [D]^2/[T]$

[b] $K_{d2} = [M]^2/[D]$

[c] $K_{d3} = [D]^2/[T^*]$

[d] $K_{d1}$–$K_{d3}$ at 298 K are shown in brackets.

**Table S3** Measured longitudinal relaxation time constants ($T_1$) for the T, T* and M peaks in A25$T^F$ $^{19}$F-NMR spectra under various temperatures.

| Peak | Temperature (K) | Conc. (μM) | Field (MHz) | $T_1$ (s) |
|---|---|---|---|---|
| T | 298 | 260 | 600 | 0.35 ± 0.01 |
| T* | 298 | 260 | 600 | 0.36 ± 0.03 |
| M | 298 | 260 | 600 | N.D.[a] |
| T | 277 | 30 | 600 | 0.34 ± 0.01 |
| T* | 277 | 30 | 600 | 0.37 ± 0.03 |
| M | 277 | 30 | 600 | 0.27 ± 0.06 |
| T | 298 | 4 | 700 | 0.31 ± 0.01 |
| T* | 298 | 4 | 700 | 0.24 ± 0.07 |
| M | 298 | 4 | 700 | 0.27 ± 0.06 |
| T | 277 | 4 | 700 | 0.31 ± 0.03 |
| T* | 277 | 4 | 700 | 0.37 ± 0.04 |
| M | 277 | 4 | 700 | 0.32 ± 0.08 |

[a] Not determined due to a low peak intensity of the M species of A25$T^F$ under this condition.

**Table S4** $^{19}$F-NMR $R_2$ for various constructs/species

| Construct (species)[a] | $^{19}$F $R_2$ ($s^{-1}$) at 277 K | $^{19}$F $R_2$ ($s^{-1}$) at 298 K |
|---|---|---|
| TTR$^F$ (T) | 57 ± 1 | 26 ± 1 |
| F87A$^F$ (M) | 34 ± 1<br>[0.60 ± 0.01] [b] | 15 ± 1<br>[0.58 ± 0.04] [b] |
| F87E$^F$ (M) | 36 ± 1<br>[0.63 ± 0.01] [b] | 15 ± 1<br>[0.58 ± 0.04] [b] |
| A25T$^F$ (T) | 62 ± 1 | 28 ± 1 |
| A25T$^F$ (T*/D) | 50 ± 2<br>[0.81 ± 0.03] [c] | 29 ± 2<br>[1.0 ± 0.1] [c] |
| A25T$^F$ (M) | 32 ± 5<br>[0.52 ± 0.08] [c] | N.D.[d] |

[a] Measured at 600 MHz with a 180°-pulse frequency of 4000 $s^{-1}$ and in NMR buffer.
[b] Relative $R_2$ ratio referenced to TTR$^F$ (T) at the same temperature.
[c] Relative $R_2$ ratio referenced to A25T$^F$ (T) at the same temperature.
[d] Not determined due to a low population of M of A25T$^F$ above 10 µM at 298 K.

**Table S5**. Activation energy determined using transition state theory and the rate constants at 277 and 298 K

| Equilibrium | T ↔ 2D | T* ↔ 2D | D ↔ 2M | T ↔ T* |
|---|---|---|---|---|
| Forward $\Delta H^{\neq}$ (kcal/mol) | $\Delta H_1^{\neq}$ =3.9 ± 0.5 | $\Delta H_5^{\neq}$ =18 ± 6 | $\Delta H_3^{\neq}$ =16 ± 7 | $\Delta H_7^{\neq}$ =-4.0 ± 1.4 |
| Forward $\Delta S^{\neq}$ (cal/mol/K) | $\Delta S_1^{\neq}$ =-40 ± 5 | $\Delta S_5^{\neq}$ =11 ± 3 | $\Delta S_3^{\neq}$ =-0.6 ± 0.3 | $\Delta S_7^{\neq}$ =-80 ± 25 |
| Reverse $\Delta H^{\neq}$ (kcal/mol) | $\Delta H_2^{\neq}$ =45 ± 4 | $\Delta H_6^{\neq}$ =56 ± 7 | $\Delta H_4^{\neq}$ =11 ± 7 | $\Delta H_8^{\neq}$ =0.02 ± 0.01 |
| Reverse $\Delta S^{\neq}$ (cal/mol/K) | $\Delta S_2^{\neq}$ =139 ± 21 | $\Delta S_6^{\neq}$=171 ± 17 | $\Delta S_4^{\neq}$ =9 ± 8 | $\Delta S_8^{\neq}$ =-61 ± 19 |

**Table S6** Comparison of thermodynamic parameters of A25T[F] with other TTR mutants published previously for the forward T↔4M equilibrium at 298 K and pH 7.0.

| Mutants | Δ*H* (kcal/mol) | TΔ*S* (kcal/mol) | Δ*G* (kcal/mol) |
|---|---|---|---|
| K80D[F, a] | -16.6 | -41.8 | 25.2 |
| K80E[F, a] | -18.0 | -44.0 | 26.0 |
| A25T[F] (T) | -33.1 | -59.0 | 25.9 |
| A25T[F] (T*) | -29.1 | -53.4 | 24.3 |
| WT[b] | N.D. | N.D. | 32.8 |

[a] Ref. 7: Also by $^{19}$F-NMR in the same NMR buffer (10 mM potassium phosphate, 100 mM KCl, pH 7.0) as this work. T = 298 K.
[b] Ref. 26: Measured by urea denaturation in 50 mM sodium phosphate, 100 mM KCl, 1 mM EDTA, 1 mM DTT at pH 7.4. T = 298 K.

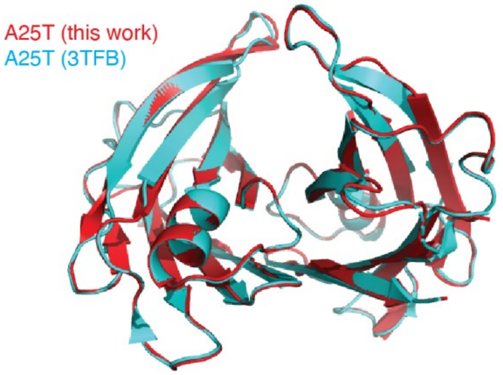


**Figure S1.** The X-ray structure of A25T from this work (red) is similar to a previously published structure (cyan, PDB 3TFB). The Cα RMSD is 0.2 Å.

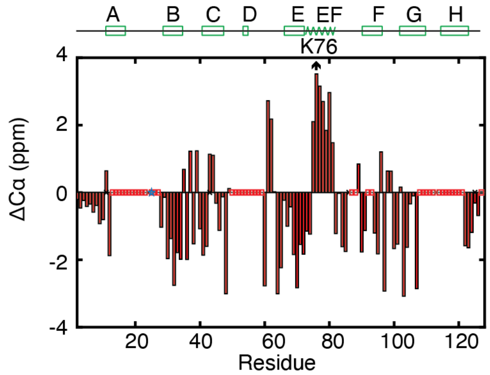


**Figure S2.** Secondary Cα chemical shift of A25T. Secondary structures taken from the A25T-structure (Figure 1A) are labeled on the top. Red boxes in the x-axis denote broadened residues in Figure 2B and the star stands for the mutated residue 25.

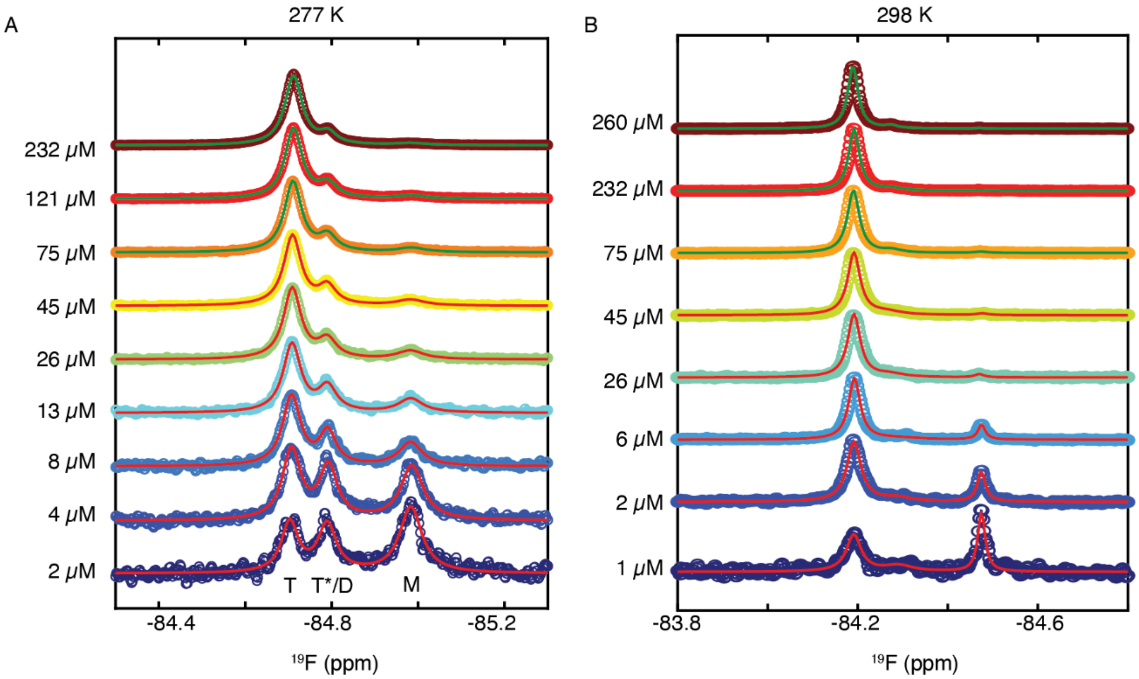


**Figure S3.** Full dilution series at 277 K (A) and 298 K (B) with $^{19}$F-NMR data of A25T$^{F}$ in circles and 3-state Lorentzian fits in solid lines.

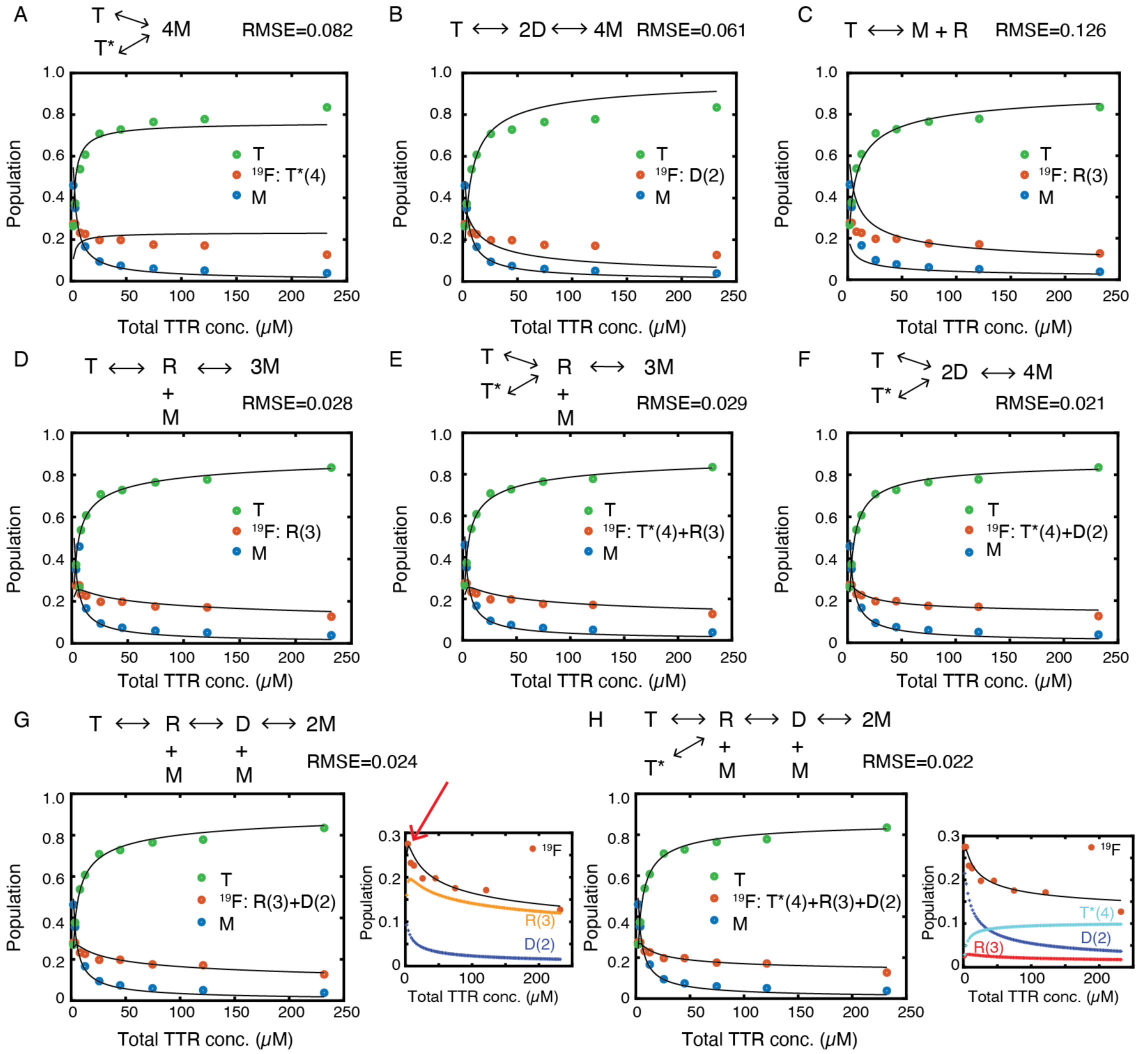


**Figure S4**. Different dissociation models to fit the A25T$^F$ dilution series at 277 K. Letters T, T*, R, D and M stand for native tetramer, mispacked tetramer, trimer, dimer and monomer. For comparison, root mean squared error (RMSE) normalized by adjusted degree of freedom is shown on the top of each panel with the associated model. The relative population of the $^{19}$F shoulder peak (red circles) is labeled with underlying species in each figure legend. Note while the T*-free sequential dissociation model (G) shows a comparable RMSE with respect to the TDM model (F), the sequential model incorrectly predicts the turnover point of the shoulder peak population around 2–4 μM (see the red arrow in the close-up view and compare with Figure 3F inset). Similar incorrect turnover points are also seen panels (D) and (E) involving trimers. The more complicated sequential model with one more fitting parameter (H) shows better consistency with low concentration data points compared to (G), but all four parameters in (H) are unconstrained. Therefore, the TDM model in (F) is the simplest model with fewest parameters constrained by the A25T$^F$ $^{19}$F-NMR data.

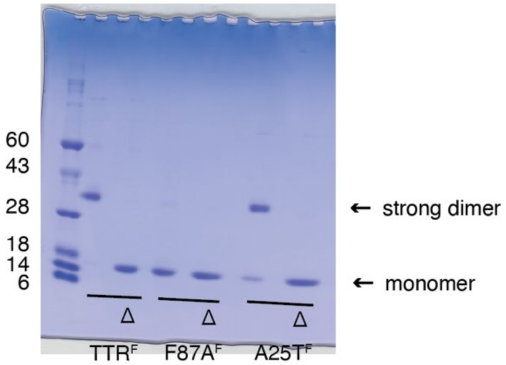


**Figure S5**. SDS PAGE analysis of boiled (right lane, marked with Δ) and unboiled (left lane) $TTR^F$, $F87A^F$ and $A25T^F$. Unboiled $TTR^F$ runs as a dimer band while boiled $TTR^F$ runs as a monomer band. Mutation of F87A at the strong dimer interface nearly abolishes the dimer band; therefore, the dimer band represents the strong dimer that is resistant to SDS-induced dissociation without boiling. $A25T^F$ without boiling maintains most of the strong dimer band, supporting that the A25T mutation noticeably perturbs but does not completely abolish the strong dimer interface as the F87A mutation.

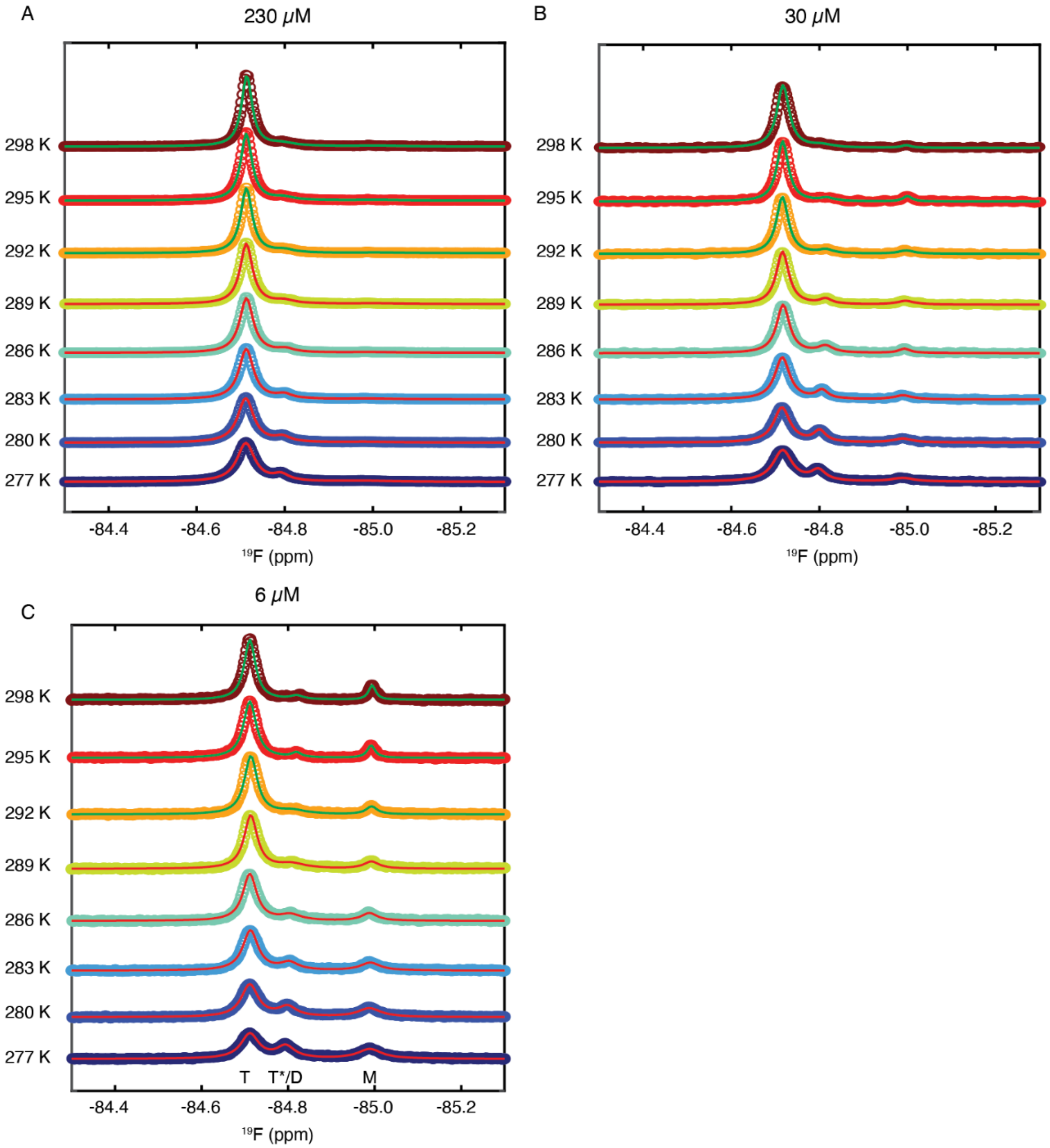


**Figure S6.** Full temperature $^{19}$F-NMR titration data sets at 230 μM (A), 30 μM (B) and 6 μM (C) of A25T$^{F}$.

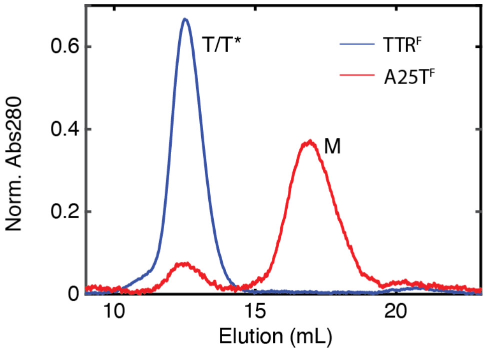


**Figure S7.** Size-exclusion elution profile of A25T$^{F}$ (4 μM) and TTR$^{F}$ (8 μM) using Superdex 75 equilibrated in NMR buffer. The absorbance at 280 nm was normalized to ensure the same total area-under-peak for both samples. The measured M population is 87% by peak area, within the range from 77% (1 standard deviation of the M peak to calculate the in-column dilution of the concentration of the M species) to 92% (2 standard deviations used in estimation) using the parameters from the van't Hoff analysis (Table 1). The elution volume corresponding to a dimeric TTR is 14.4 mL where the absorbance at 280 nm is low.

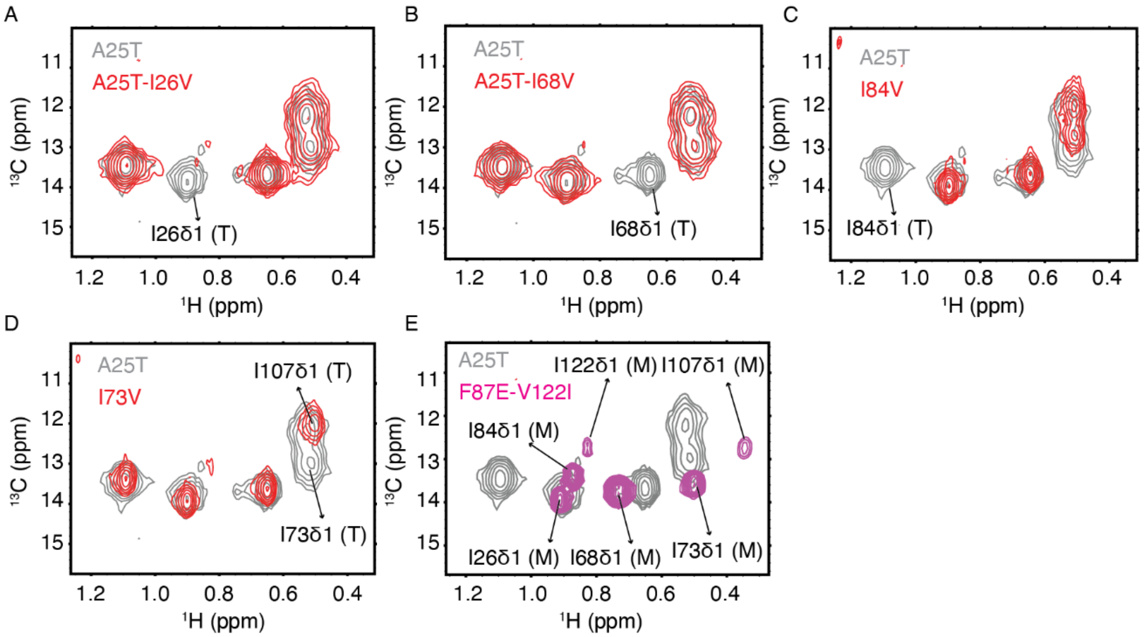


**Figure S8**. Assignment of Ile δ1 methyl resonances by mutagenesis. The $^{13}$C HMQC spectrum of A25T at 298 K (gray) overlaid with spectra of A25T-I26V (red, in A), A25T-I68V (red, in B), WT-I84V (red, in C), WT-I73V (red, in D) and a monomeric F87E-V122I (magenta, in E). Letters T and M denote tetramer and monomer, respectively. The WT-I73V construct was used because the A25T-I73V construct was insoluble. The WT-I84V construct was used as the yield of A25T-I84V was very low. The I68δ1 and I84δ1 cross peaks in the F87E-V122I spectrum in (E) overlap with the two minor peaks in A25T respectively.

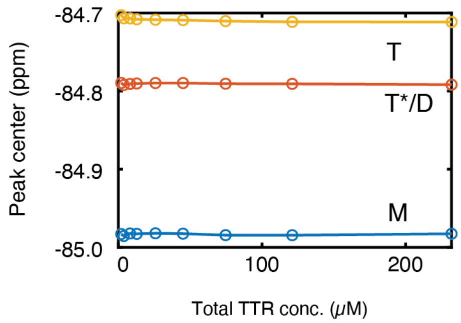


**Figure S9.** The $^{19}$F chemical shifts for the M, T*/D and T states of A25T$^{F}$ are concentration-independent at 277 K.

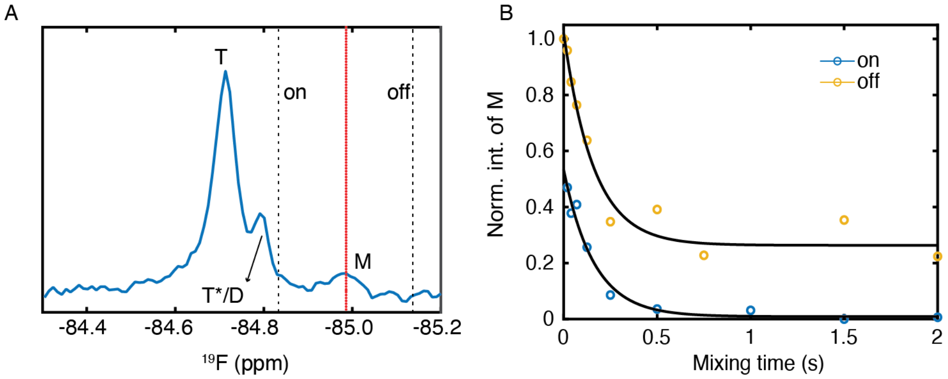


**Figure S10.** $^{19}$F saturation transfer experiment to measure M → D exchange rate for 30 μM A25T$^F$ at 277 K. (A) $^{19}$F-NMR spectrum of A25T$^F$ with labeled on and off resonance offsets (dashed line) as well as the M peak center (red dotted line). Although T* and D have degenerate $^{19}$F chemical shifts under the shoulder peak, only D is directly connected with M in the TDM scheme. (B) M peak intensity fitted by single exponential functions for both on and off resonance experiments. The M → D rate calculated by the intensity plateau difference using Eq. 9 is 1.3 ± 0.3 s$^{-1}$. This rate is within error of the rate (2.5 ± 1.2 s$^{-1}$) extracted by the single exponential fits, confirming the accuracy of the measurements.

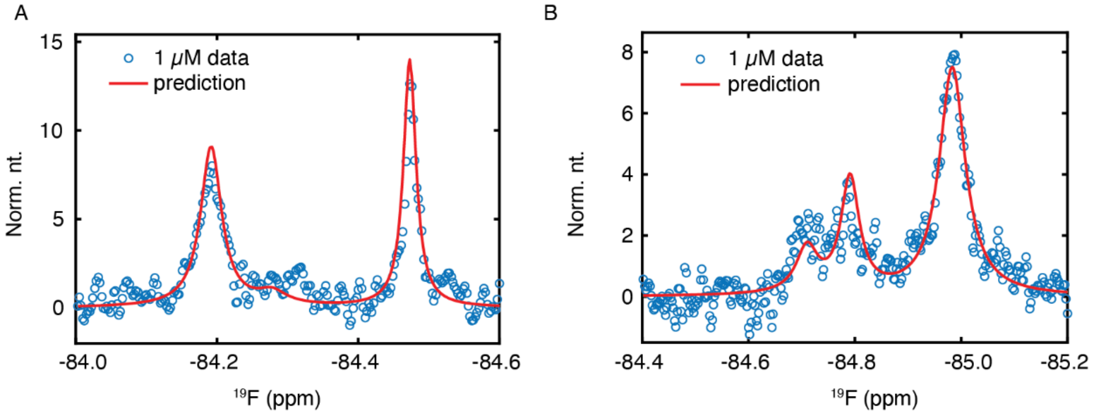


**Figure S11**. Comparison between measured data (blue circles) of 1 μM A25T$^{F}$ at 298 K (A) / 277 K (B) and line shapes predicted from fitting results in Tables 2 and 3. Due to low signal/noise ratio, the 1-μM data were not used in the line shape fitting. Nevertheless, the spectral comparisons indicate that the line shape fitting using data greater than 1 μM reasonably recapitulates main features in the 1-μM data at both temperatures.

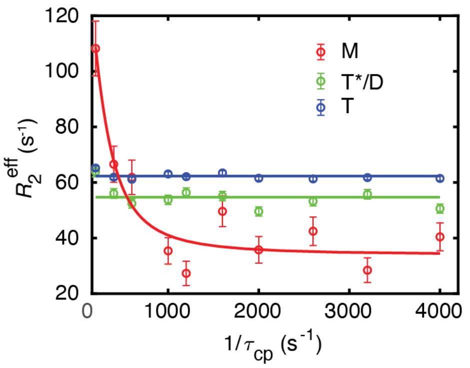


**Figure S12**. CPMG fit of the effective $^{19}$F $R_2$ of the A25T$^F$ M peak at 30 μM and 277 K. The solid line denotes the fast exchange Luz-Meiboom fit for the M $^{19}$F $R_2$ data with $k_{ex}$ = 780 ± 320 s$^{-1}$ and $R_{2,0}$ = 34 ± 3 s$^{-1}$. For comparison, the $^{19}$F $R_2$ data for T* and T peaks, which exhibit no relaxation dispersion, are plotted with flat lines that are obtained from averaging all respective $^{19}$F $R_2$ data points.

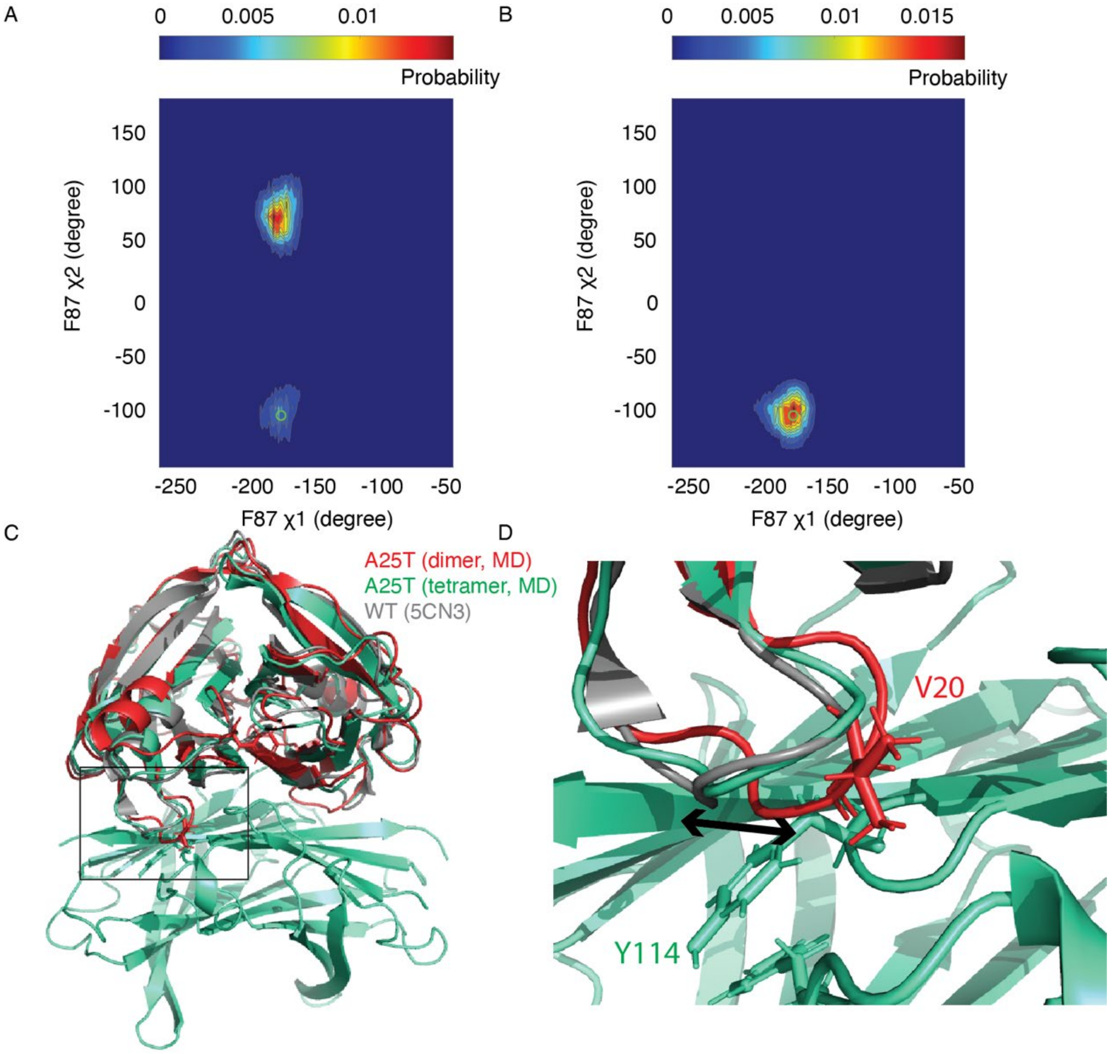


**Figure S13.** (A–B) 2D probability contour plots for the A25T dimer (A) and tetramer (B), extracted from trajectories shown in Figures 9A–B. The green circle marks the averaged χ1 and χ2 of the F87 side chain from the two chains in our A25T X-ray structure. (C) Structural comparison showing the perturbation of the AB loop in simulated A25T MD structures compared to these from A25T tetramer MD simulations or the WT TTR X-ray structures (PDB 5CN3). The A25T dimer MD snapshot is taken from Figure 9C. (D) Close-up view of the AB loop in one A25T protomer in the simulated A25T dimer clashes with a neighboring protomer in a simulated A25T tetramer that is aligned in (C). By contrast, the AB loops in the WT X-ray and A25T tetramer MD structures take on alternative conformations that do not have steric hindrance for the packing around the weak dimer interface.

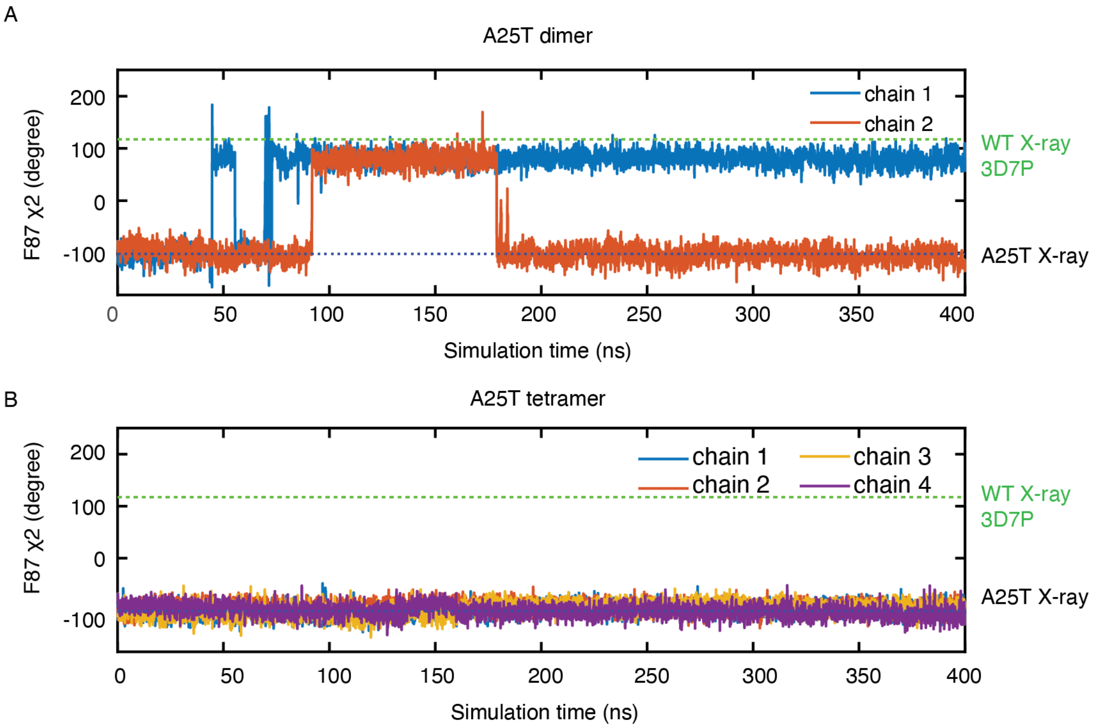


**Figure S14.** (A–B) F87 side chain dihedral χ2 dynamics for A25T dimer (A) and tetramer (B) from canonical ensemble ensemble simulations. F87 χ2 from the A25T X-ray structure (black) and a low pH structure of WT (green, PDB 3D7P) is plotted for reference.
